# Electrostatic properties of disordered regions control transcription factor search and pioneer activity

**DOI:** 10.1101/2025.03.07.641980

**Authors:** Sim Sakong, Beat Fierz, David Suter

## Abstract

Transcription factors (TFs) recognize specific target sites to control gene expression. While prokaryotic TFs combine 1D and 3D diffusion to efficiently locate their binding sites, the target search of eukaryotic TFs within the chromatinized genome is poorly characterized. Here we combine *in vivo* and *in vitro* single-molecule imaging to dissect the role of DBD-flanking regions (DFRs) of the mammalian Sox2 and Sox17 TFs in searching for their binding sites. We demonstrate that the DFR_Sox2_ dramatically enhances the search efficiency of Sox2 as compared to the negatively-charged DFR_Sox17_. An enhanced search on naked DNA is primarily mediated by an increase in target recognition rate during 1D sliding. In contrast, target site recognition within nucleosomal DNA is mainly enhanced by increased nonspecific interactions with nucleosomes, facilitating binding to closed chromatin and reinforcing pioneering activity. These findings provide critical insights into the biophysical mechanisms governing TF target search on the chromatinized genome.

## INTRODUCTION

TFs are central regulators of gene expression in all living organisms. They identify and bind to target DNA sites in a sequence-specific manner via their DBDs (specific interactions), navigating a vast landscape of nonspecific DNA (nonspecific interactions). Efficient target search is crucial for achieving a sufficiently high association constant, especially given the limited number of TFs available within the compact, fractal-like nuclear environment. TF copy numbers vary widely, ranging from 10^3^ to 10^6^ molecules per cell^1–3^. Collectively, these molecules need to cover tens of thousands TF target sites identified by chromatin immunoprecipitation sequencing (ChIP-seq)^2,4,5^. These sites typically correspond to regulatory elements such as enhancers and promoters. However, the number of potential binding motifs across the genome is in the range of millions. Therefore, TFs face the challenge of efficiently locating their specific binding sites while frequently encountering numerous nonspecific decoy sites along their search path^6–10^.

In particular within chromatin, the mechanisms by which TFs maximize target search efficiency remain poorly understood. The facilitated diffusion model, formulated for free DNA, proposes that TFs enhance search efficiency by leveraging nonspecific DNA interactions, alternating between 1D diffusion along DNA and 3D diffusion within the nucleoplasm^11–14^. This dimensionality-reduced diffusion process increases search efficiency by orders of magnitude as compared to free diffusion alone. Earlier studies have shown that LacI follows the facilitated diffusion model in live E. coli^15^ and *in vitro*^16–18^. However, in eukaryotes, chromatin acts as a barrier to TF-DNA interactions, potentially impairing TF target search and site recognition^19–22^. A subset of eukaryotic TFs, known as pioneer TFs (pTFs), thus possess the ability to bind nucleosomal DNA and initiate local chromatin opening, facilitating access to previously inaccessible regulatory elements^23–26^.

Eukaryotic TFs contain intrinsically disordered regions (IDRs) outside of their DBDs. However, how these regions influence the target search process remains unclear. Recent findings indicate that IDRs play an important role in defining the binding specificity of TFs *in vivo*^27^. TFs with identical DBDs but distinct IDRs recognize the same motifs *in vitro*, but they often bind distinct genomic subsets *in vivo*, highlighting the role of DBD-proximal IDRs in modulating target selection. Mechanistically, IDRs, in particular the trans-activation domains, are known to mediate protein-protein interactions^28–31^. However, even in the absence of direct interaction partners, DBD-flanking IDRs (DFRs) may play a key role in directly defining chromatin binding and diffusion behavior^32–35^.

Sox TFs bind their targets through their high mobility group box DBD, either alone or as heterodimers with other TFs. Sox2 is a core pluripotency TF heterodimerizing with Oct4 and is well-characterized for its potent pioneer activity^5,25,36^. Another member, Sox17, can also heterodimerize with Oct4 and is expressed during differentiation towards extraembryonic endoderm. While Sox2 and Sox17 alone bind to a virtually identical motif^37^, they differ in the composite motif that they bind with Oct4 due to differences in their heterodimerization interface, located within the DBD^38,39^. Sox2 and Sox17 were also shown to markedly differ in their ability to colocalize with mitotic chromosomes, with Sox2 exhibiting strong binding, whereas Sox17 associates only weakly^40^. The strong mitotic chromosome association of Sox2 has been proposed to reflect its superior ability to nonspecifically bind chromatinized DNA and to efficiently search the genome. The Sox2 domain responsible for this property and its role in Sox2’s potent pioneer activity remain unclear.

Here, we investigated the role of differentially-charged DFR stretches of the Sox2 and Sox17 TFs (DFR_Sox2_ and DFR_Sox17_, respectively) in regulating their ability to search for their binding sites on naked DNA and chromatin, using both *in vivo* and *in vitro* single-molecule approaches.

## RESULTS

### Sox2 and Sox17 mutants to study the DFR impact on target search

Sox2 and Sox17 differ in amino acid composition of the region C-terminal to the DBD, being more negatively-charged in Sox17 compared to Sox2 **(Figure 1A)**. We thus hypothesized that the DFR, by virtue of its distinct charge distribution, is critical for the highly efficient target search and the strong pioneer activity of Sox2. To test this hypothesis, we determined the impact of swapping DFRs between the two factors. We defined the DFRs as a roughly 100 amino acid-long stretch, C-terminal of the DBD with overall charge DFR_Sox2_ (+1 z) and DFR_Sox17_ (−9.5 z) **(Figure S1A)**. We then generated a pair of DFR-swapped constructs: i) Sox2 containing the acidic DFR_Sox17_ (Sox2a) and ii) Sox17 containing the DFR_Sox2_ (Sox17b) **(Figure 1B)**. We also generated constructs containing only the DBDs of Sox2 and Sox17 (Sox2D and Sox17D) to eliminate the impact of DFRs on Sox-DNA binding dynamics.

**Figure 1.**
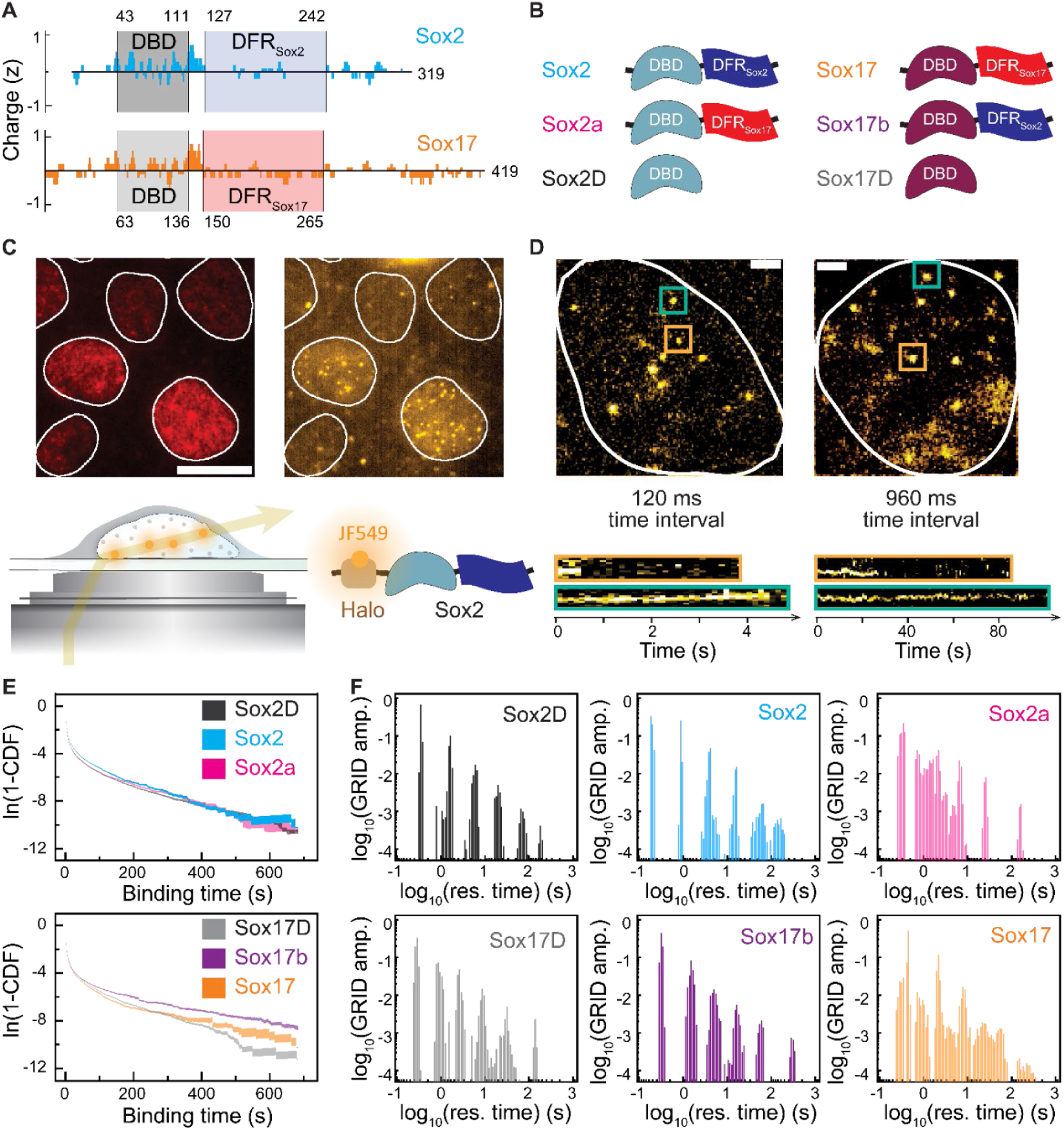
DFR charge controls the distribution of TF residence times. (A) Charge distributions along the sequence of Sox2 (top) and Sox17 (bottom). (B) Scheme representation of the Sox TF library. (C) Top: Representative images of in vivo SMT. Nuclei (left, red) and sparsely labeled Sox TFs (right, yellow) are visualized in the SiR647 and JF549 emission channels, respectively. Scale bar: 15 μm. Bottom: Scheme of *in vivo* single-molecule imaging using HILO microscopy (left) and a labeling strategy for Sox TFs using the Halo-JF549 dye (right). (D) Representative images of JF549-labeled Halo-Sox2 molecules, acquired at different time intervals in time-lapse movies. Scale bar: 2 μm. The kymographs below each image correspond to the molecules in the highlighted squares. (E) Cumulative distribution of Sox TF binding times. Curve shadings: standard deviation (SD) calculated from 100 resampling iterations, each using 80% of the dataset selected randomly (N = 281, 169, 223, 91, 164, 159 cells). (F) GRID spectra of Sox TF residence times. The GRID amplitude indicates the bound fraction per unit of time for the corresponding residence times. The spectra were generated from the same resampled data in (E).

To investigate the impact of the DFRs on the target search process in living cells, we engineered mESC lines (based on the CGR8 cell line) containing HaloTag (Halo) fusions to either Sox2, Sox2a, Sox17, Sox17b, Sox2D or Sox17D stably integrated under the control of a doxycycline (dox)-inducible promoter. This allowed us to express Sox TFs at low levels, thereby minimizing their phenotypic impact and facilitating single-molecule imaging^41^ **(Figure 1C)**. We then characterized the genome-wide binding profiles of this set of TFs via ChIP-seq. Motif enrichment analysis of ChIP-seq peaks indicated that all Sox TFs bound to nearly identical motifs, including their respective canonical motifs for Sox2 and Sox17 **(Figure S1B)**. Therefore, although Sox17 is not expressed endogenously in mESCs, Sox17 and its mutant Sox17b can still recognize their target sites in this context. Notably, Sox2a bound to motifs that are more similar to those bound by Sox2 than Sox17, confirming that the heterodimerization properties of Sox2 are independent of its DFR.

Since Sox2 is crucial for maintaining pluripotency in mESCs, we also determined whether Sox2a can fulfill this role. To this end, we performed a pluripotency rescue assay by using the 2TS22C cell line^42^, which allows to fully deplete Sox2 upon dox addition within 48 hours. These cells were also engineered to express Halo-Sox2 or Halo-Sox2a at the same level. We then treated these cell lines with dox for 5 days followed by quantification of the percentage of undifferentiated cell colonies and of NANOG expression levels via immunofluorescence (IF) **(Figures S1C-D)**. We found that Halo-Sox2a was able to rescue ESC colony morphology and NANOG expression, even though less potently than Halo-Sox2, demonstrating that Sox2a can partially replace Sox2 in maintaining pluripotency of mESCs.

### DFRs impact the residence times of TFs on DNA *in vivo*

Single-molecule tracking (SMT) allows direct observation of the TF search process within living cells. We expressed our six different TF constructs at low level and labeled them with JF549^43^. To label chromatin, we used SNAP-H2B labeled with silicon rhodamine (SiR)^44^. We then quantified TF-DNA interaction dynamics using single-molecule highly inclined and laminated optical sheet (smHILO) microscopy. We first used four different imaging conditions by maintaining 60 ms of illumination time and varying dark time to obtain imaging intervals of 120, 240, 480 and 960 ms. This approach allows to determine TF binding behavior over a broad temporal range **(Figure 1D)**. We observed that most individual molecules were confined within nuclei **(Figures 1C-D)**. We then reconstructed trajectories for individual molecules using TrackIt^45^ and plotted the reverse cumulative density function (1-CDF) of binding times. Sox2 and Sox17b exhibited slower 1-CDF decays, compared to Sox2a and Sox17 or their DBD alone **(Figures 1E, S1E)**, indicating slightly longer residence times.

The TF bound states are influenced by various factors, such as chromatin accessibility, RNA binding, and protein-protein interactions^37,46–48^. These diverse subpopulations are expected to exceed the binary classification of specific versus nonspecific bound states^2,49,50^. To resolve populations of bound molecules in an unbiased manner, we applied the genuine rate identification (GRID) method^47^, which uses an inverse Laplace transform of the 1-CDF to infer the underlying distribution of residence times. The resulting GRID spectrum represents the relative fractions and corresponding residence times of dissociation processes for Sox-DNA complexes throughout the observation period **(Figure 1F)**.

For Sox2 and Sox17b as well as Sox2D and Sox17D, we identified six well-resolved populations of bound states from a distribution of residence times ranging from 0.1 s to 1000 s. Conversely, Sox2a and Sox17 exhibited broad residence time distributions, suggesting complex interactions associated with a continuum of binding energies. The amplitudes, corresponding to state populations, decreased for increasing residence times, and in particular the population associated with long binding events (>10 s) was increased for Sox2 (0.042) vs Sox17 (0.018) **(Figure S1F; Table S1)**. In contrast, the short-lived and long-lived populations for Sox17 vs Sox17b and Sox17D vs Sox2D were comparable. Overall, these results suggest that DFRs have an impact on Sox TF-DNA residence times.

### DFRs regulate *in vivo* target search efficiency

Next, we dissected the target search process of the different Sox TFs. To quantify search efficiency, we first determined *in vivo* association rates through diffusion analysis. We performed continuous imaging at 10 ms acquisition to observe both diffusing and bound molecules **(Figure 2A)**. As previously reported for Sox2^51,52^, we assumed that all Sox TFs adhere to a three-state diffusion model, encompassing fast diffusion, slow diffusion, and a bound state **(Figure 2B)**. To verify this assumption, we constructed 1-CDFs of jump distances for each Sox TF, which were well described with a triexponential function, representing three diffusing populations **(Figure S2A)**.

**Figure 2.**
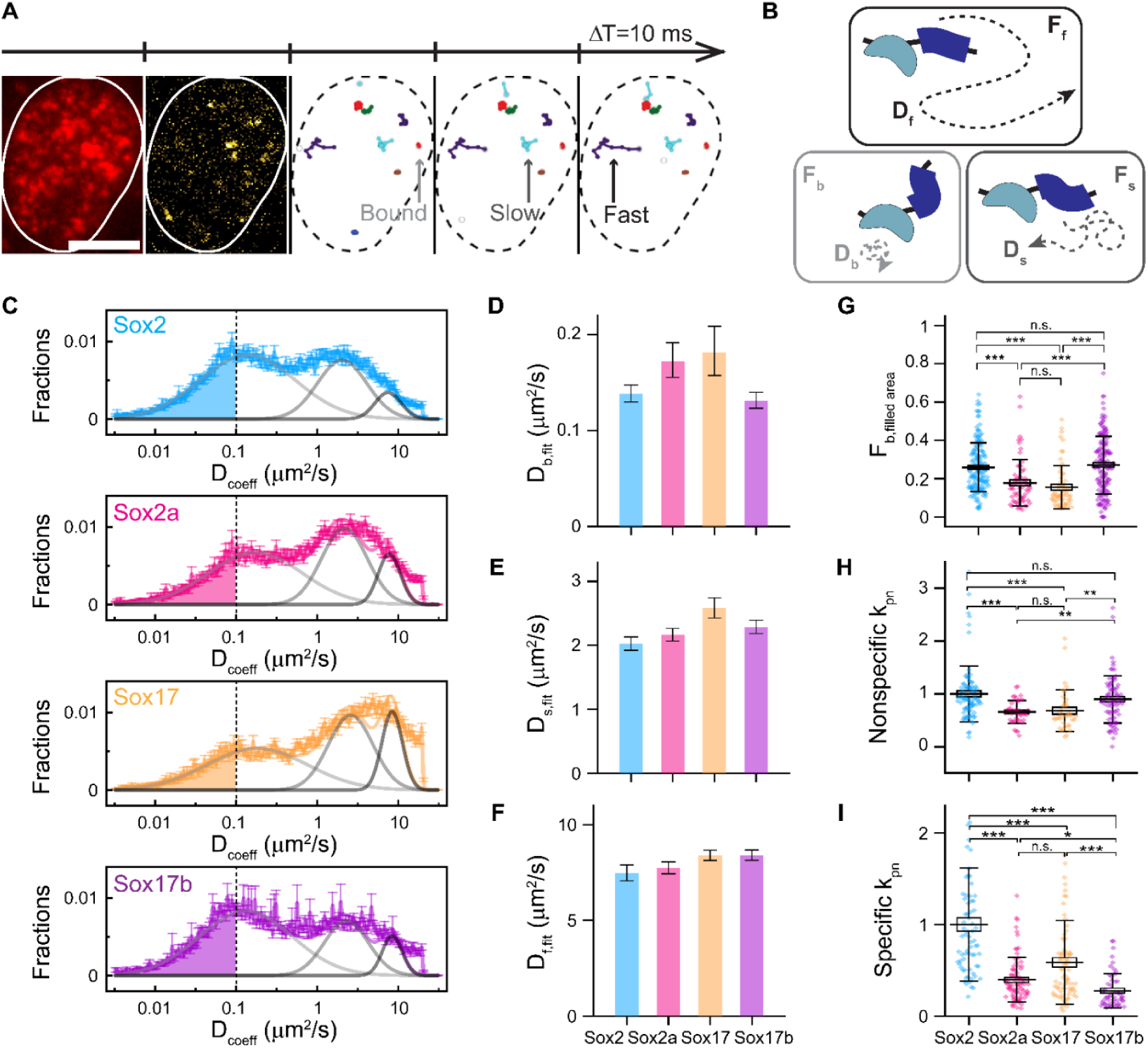
DFR charge controls DNA binding affinity and target search efficiency. **(A)** Representative images from a movie showing H2B-SNAP (red) and Halo-Sox2 (yellow), with color-coded trajectories of Sox2 molecules. Scale bar: 5 μm. **(B)** Scheme of the three-state diffusion model with kinetic parameters for diffusion coefficients (D) and corresponding fractions (F) for fast (f), bound (b), and slow (s) diffusion modes. **(C)** Diffusion coefficient distributions of Sox TFs molecules based on single-cell analysis (N = 86, 38, 38, 72 cells). The distributions were fitted (solid color) with three Gaussian functions (gray-scaled). **(D-F)** Peak diffusion coefficients of the three Gaussian fits shown in (C) of bound mode (light gray) **(D)**, slow diffusion (dark gray) **(E)**, and fast diffusion (black) **(F)**. Bar: fitting values. Cap: fitting error. (G) Bound fractions of molecules with D_coeff_<0.1 μm^2^/s (colored area) shown in (C). Line: mean. Cap: SD. Box: standard error of mean (SEM). **(H-I)** Pseudo on-rates of nonspecific (N = 70, 81, 78, 54 cells) **(H)** and specific (N = 86, 40, 38, 80 cells) **(I)** interactions per cell, normalized to the mean of Sox2. Line: mean. Cap: SD. Box: SEM. Statistical significance was evaluated using Dunn-test following a nonparametric Kruskal-Wallis ANOVA (*: p≤0.05; **: p≤0.01; ***: p≤0.001).

We then computed the diffusion coefficients from the trajectories and plotted the distribution across different Sox TFs **(Figure 2C**; see **Methods)**. The distributions were fitted with three different gaussian functions representing the three diffusing modes **(Figures 2D-F; Table S1)**, revealing no significant differences between the diffusion coefficients for all Sox TFs.

To quantify the bound fraction of each Sox TF, we defined molecules with a diffusion coefficient <0.1 μm^2^/s as immobile^32,53,54^. Of note, this bound mode encompasses all the GRID bound states (see also **Methods)**. This revealed that Sox TFs with a more positively-charged DFR, Sox2 and Sox17b, showed higher bound fractions than Sox2a and Sox17 **(Figure 2G)**. To orthogonally confirm the kinetic parameters obtained from the diffusion coefficient distributions, we extracted them from the jump distance distributions using Spot-On^51^. This method additionally applies lateral movement correction accounting for particles moving out of the focal plane. Consistent with previous analysis **(Figure 2G)**, Sox2 and Sox17b exhibited higher bound fractions than Sox2a and Sox17 **(Figure S2B; Table S1)**. Notably, the slow-diffusing populations of Sox2 and Sox17b peaked at fewer jumps than those of Sox2a and Sox17 **(Figure S2C)**, suggesting that they may locate their binding sites more rapidly during slow diffusion.

Next, we investigated whether the set of TFs differ in their association rate to genomic targets. We measured their pseudo on-rate (*k_pn_*)^40^, which is defined as the time-averaged binding frequency of TFs to the whole genome, as determined by the number of immobilized molecules per imaging frame (see **Methods)**. We defined 1 s as a minimal trajectory length to separate nonspecific and specific interactions to determine *k_pn_* **(Figures 2H-I; Table S1)**. Sox2 (*k_ns,pn_*=1.00±0.53) and Sox17b (*k_ns,pn_*=0.89±0.45) displayed higher nonspecific *k_pn_* than Sox2a (*k_ns,pn_*=0.66±0.22) and Sox17 (*k_ns,pn_*=0.68±0.39) **(Figure 2H)**, in line with previous diffusion analysis, which suggests that DFR_Sox2_ leads to a high frequency of nonspecific DNA binding events for Sox2 and Sox17b. Importantly, Sox2 (*k_s,pn_*=1.00±0.62) exhibits a faster specific *k_pn_* than all the other TFs (*k_s,pn_*=0.40±0.24 for Sox2a, 0.59±0.46 for Sox17, 0.28±0.19 for Sox17b) **(Figure 2I; Table S1)**. This suggests that DFR_Sox2_ is necessary but not sufficient to ensure a high specific *k_pn_* of Sox TFs. In summary, our *in vivo* data demonstrate that TFs containing DFR_Sox2_ interact more frequently and nonspecifically with DNA and display enhanced target search efficiency compared to TFs containing DFR_Sox17_.

### DFRs do not impact 3D diffusion-mediated association to naked DNA and residence times *in vitro*

A limitation of *in vivo* measurements is that specific and nonspecific binding events can only be inferred from residence times. Furthermore, the *in vivo* chromatin context in which these events occur cannot be identified. We thus performed *in vitro* studies to examine how DFRs impact DNA binding dynamics of Sox TFs on naked and chromatinized DNA, focusing solely on Sox2 and Sox2a. We employed a fluorescence colocalization approach^24^ using single-molecule total internal reflection fluorescence (smTIRF) microscopy. In this assay, Alexa Fluor 647 (AF647)-labeled DNA or chromatin substrates are immobilized in a flow channel and their position is detected in the AF647 channel **(Figure 3A)**. Subsequently, Sox TFs are introduced, which carry a JF549-labeled Halo for detection. When bound to DNA, they appear as fluorescent spots colocalizing with the immobilized DNA molecules. Recording movies enabled us to directly determine on-rates and residence times for specific DNA or chromatin environments.

**Figure 3.**
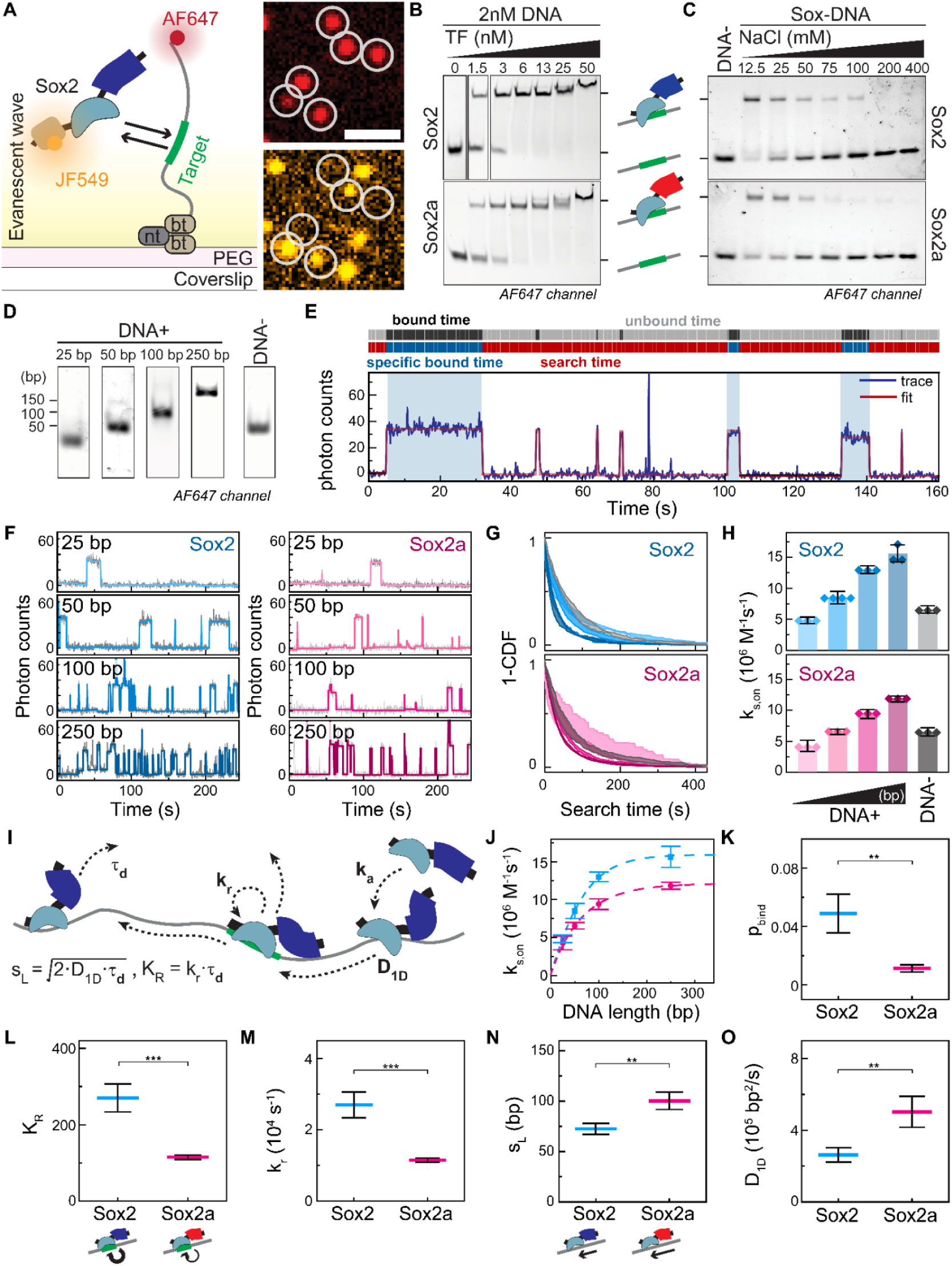
DFR charge controls target recognition efficiency on naked DNA and slow 1D diffusion. **(A)** Left: Scheme of single-molecule experiment to detect Sox TFs binding to DNA. bt: biotin, nt: neutravidin. Right: Microscopy images showing DNA loci in the AF647 emission channel (top, red), and bound Sox2 in the JF549 emission channel (bottom, yellow). Scale bar: 2 μm. **(B)** EMSA of indicated TFs. **(C)** EMSA at different NaCl concentrations for indicated TFs. **(D)** DNA templates of different lengths used for single-molecule binding experiments. Fluorescence time-trace of Sox2 binding to 50 bp motif-containing DNA, detected by JF549 emission (blue) and fitted with a step-detection algorithm (red). **(F)** Fluorescence time-traces of Sox TFs binding to DNA of indicated lengths. The raw data (gray) were fitted (blue, red). **(G)** Cumulative distributions of search times for Sox TFs, on various DNA substrates. Curve shading: SD (N=3-4). **(H)** Specific on-rate of Sox TFs obtained by fitting 1-CDF curves in (G) with a monoexponential decay. Symbols: replicates (N=3-4). Bar: mean. Cap: SD. **(I)** Scheme of the 1D target search model (for details see text). **(J)** Specific on-rate curve as a function of DNA length. Fit: 1D target search model. Symbols: mean. Cap: SD. **(K-O)** Fitting and derived parameters from the fitting in (J) (see Methods): target recognition probability during 1D sliding **(K)**, target recognition constant **(L)**, target recognition rate **(M)**, sliding length **(N)**, and 1D diffusion coefficient **(O)**. Line: mean. Cap: fitting error. For the statistical significance, two-tailed t-tests were performed (*: p≤0.05; **: p≤0.01; ***: p≤0.001).

We produced recombinant Halo-containing Sox2 and Sox2a in insect cells **(Figure S3A;** see **Methods)**. We validated the ability of Sox2 and Sox2a to bind their specific sites using electrophoretic mobility shift assays (EMSA), which allowed us to estimate dissociation constants of 1.6 nM for Sox2 and 2.3 nM for Sox2a **(Figure 3B)**, indicating comparable binding affinities within the reported range for Sox2 (1.50-3.83 nM)^55^. We also measured the salt dependency of Sox-DNA binding via EMSA by varying NaCl concentrations. Sox2a dissociated from DNA at lower NaCl concentrations (EC_50,Sox2a_=27.1±6.2 mM NaCl) compared to Sox2 (EC_50,Sox2_=38.1±1.8 mM NaCl) **(Figures 3C, S3B-C)**. Therefore, Sox2 is able to bind its consensus site at higher ionic strength than Sox2a, suggesting a charge-dependent effect of DFRs on the binding mechanism.

We then investigated how the charged DFRs influence TF target search on naked DNA. We designed DNA substrates of 25, 50, 100 and 250 bp length, based on the 601 Widom sequence (601WS)^56^ **(Figure 3D)**. Each DNA substrate contained a single Sox motif (5’- CTTTGTT-3’) at the center (DNA+), except for a 50 bp control DNA devoid of the motif (DNA-). We labeled the DNA duplexes with AF647, attached a biotin moiety to the 5’ end, immobilized them in a flow cell and determined their position by smTIRF imaging. We then injected Sox TFs at a concentration of 2 nM into the cell, and recorded 10-minute movies in the JF549 channel at a frame rate of 5 Hz with a 100 ms illumination time to monitor Sox-DNA binding events by colocalization detection **(Figure 3A)**. To analyze the binding kinetics, we extracted fluorescence time-traces of the JF549 channel, where high intensity reflected bound states of the Sox TFs **(Figure 3E)**. This revealed that the frequency of binding events for both Sox2 and Sox2a increased with longer DNA, in particular for short binding events **(Figure 3F)**.

We next determined the on-rates (*k_on_*) for Sox2 and Sox2a. To this end, we analyzed the 1-CDF of unbound times **(Figure S3D)** via fitting with a monoexponential decay. This revealed that Sox2 and Sox2a displayed comparable *k_on_* values across all DNA substrates **(Figure S3E; Table S2)** with *k_on_* increasing with DNA length. Moreover, *k_on_* of DNA− was similar to *k_on_* of 50 bp DNA+. This confirms that DNA length, not the presence of a specific motif, determines the overall frequency of TF-DNA interactions.

To determine the intrinsic ability of a TF to associate with DNA, we eliminated the impact of DNA length by defining the nanoscopic association rate (*k_a_*) as the association rate to any single site on DNA (specific or nonspecific). We observed that *k_on_* exhibited a nonlinear relationship with DNA length in bp **(Figure S3F)**. The collision radius of two molecules in solution is proportional to their radius of gyration, and this becomes particularly important for DNA lengths above the DNA persistence length of 45 nm^57–59^. Indeed, *k_on_* scaled directly with the calculated DNA gyration radius **(Figure S3G;** see **Methods)**. Correcting for this effect, we found values of *k_a,Sox2_* = 2.13±0.13×10^5^ M^−1^s^−1^bp^−1^ and *k_a,Sox2a_* = 1.91±0.24×10^5^ M^−1^s^−1^bp^−1^ **(Figure S3H)**. This indicates that DFRs do not impact the overall probability of interactions between freely diffusing TFs and naked DNA.

We then quantified TF residence times by analyzing the distribution of bound times. The 1-CDF curves were well described by a biexponential decay **(Figure S3I)**, matching two binding modes of Sox2 and Sox2a with DNA+ **(Figure S3J; Table S2)**: a short-lived (*τ_1,Sox2_*=1.1±0.3 s, *τ_1,Sox2a_*=0.8±0.3 s) and a long-lived interaction mode (*τ_2,Sox2_*=14.9±4.5 s, *τ_2,Sox2a_*=12.4±5.2 s). Conversely, the binding of Sox2 and Sox2a with DNA− was characterized by a single short-lived population (*τ_2,Sox2_*=0.5±0.01 s, *τ_2,Sox2a_*=0.6±0.05 s), indicating that the long-lived population corresponds to target binding. The average residence times and bound fractions for all DNA lengths and both TFs **(Figures S3J-K)** aligned well with a previously reported value for Sox2 measured by *in vitro* SMT^60^. For each DNA length, we observed similar specific residence times *τ_2_* for Sox2 and Sox2a **(Figure S3L)**. In contrast, it was not possible to directly relate *τ_1_* to nonspecific interactions. Previous measurements using fluorescence correlation spectroscopy in mouse embryos revealed that such nonspecific DNA interactions at individual random sites are extremely short-lived, with microscopic nonspecific residence times *τ_d_*=10 ms^61^. Similarly, the nonspecific residence time of LacI in E. coli has been reported as less than *τ_d_*=5 ms using SMT^62^. These times are well below the time resolution of our experiment (5 Hz) **(Figure S3L)**. This suggests that the short-lived interactions we measured as *τ_1_* likely represent the extreme tail of the nonspecific residence time distribution, and may involve numerous transient events of Sox TFs, such as hopping^11,17^, where TFs repeatedly re-associate and dissociate within a local regime before fully dissociating.

### DFR_Sox2_ facilitates target search by increasing the target recognition probability

The overall DNA binding rate (*k_on_*) is determined by the intervals between all binding events, regardless of interaction type. Because it encompasses both specific and nonspecific interactions, *k_on_* does not directly reflect the efficiency in locating a specific target site. Target search efficiency, in contrast, is defined solely by the intervals between specific interactions, which we refer to as ‘search time’.

To quantify target search efficiency, we introduce a new rate constant: the specific on-rate (*k_s,on_*). To determine *k_s,on_*, we used a residence-time threshold of *τ_res_*>1 s to identify specific binding events and determined the search times (as illustrated in **Figure 3E)**. We fitted the search time distributions **(Figure 3G)** with a monoexponential decay to determine *k_s,on_* for Sox2 and Sox2a interacting with the different DNA substrates. This analysis revealed that *k_s,on_* increased with DNA length for both Sox2 and Sox2a **(Figure 3H; Table S2).** We interpret this increased search efficiency for both Sox TFs on longer DNA as evidence that 1D sliding facilitates target search^11^. Sox2 consistently exhibited about 1.5 times higher *k_s,on_* values compared to Sox2a, suggesting that DFR_Sox2_ enhances target search efficiency over DFR_Sox17_.

We then implemented a theoretical formulation of the ‘facilitated diffusion model’ describing the target search of a TF on DNA containing a single target site^11,15^ **(Figure 3I)**. Here, the target search process involves three steps. First, a TF binds nonspecifically to a random DNA site. Second, it slides along the DNA to the target until it either dissociates, or recognizes and specifically binds the target site. The model is thus defined by four parameters: i) the nanoscopic association rate (*k_a_*), previously determined as about 2×10^5^ M^−1^s^−1^bp^−1^ for both Sox2 and Sox2a; ii) the 1D diffusion coefficient on DNA (*D_1D_*); iii) the target recognition rate (*k_r_*), which determines the probability of the TF recognizing the specific DNA site when sliding over it; and iv) the nonspecific residence time *τ_d_*, reporting on the probability of the TF dissociating from nonspecific DNA sites, and set to 10 ms for both TFs^61^. Since *τ*_d_ sets the intrinsic timescale of the search process, we defined both the sliding and target recognition processes as a function of *τ_d_*, introducing an average sliding length (*s_L_*^2^ = 2·*D_1D_* · *τ_d_*) and a target recognition constant (*K_R_* = *k_r_* · *τ_d_*) to reduce the number of parameters **(Figure 3I**; see **Methods)**.

The 1D target search model, now a function of two free parameters, *K_R_* and *s_L_*, was subsequently applied to fit the *k_s,on_* dependence on DNA length **(Figure 3J; Table S3)**. Computing the probability that TFs bind to the target site specifically during 1D sliding (*p_bind_*) resulted in *p_bind,Sox2_*=0.049 and *p_bind,Sox2a_*=0.011 **(Figure 3K)**, suggesting that the DFR_Sox2_ facilitates the TF target search. Accordingly, *K_R_*, which was determined to be 2.3-fold higher for Sox2 **(Figure 3L)**, resulted in a higher target recognition rate for Sox2 (*k_r,_*_Sox2_=2.70±0.36×10^4^ s^−1^) over Sox2a (*k_r_,*_Sox2a_=1.15±0.06×10^4^ s^−1^) **(Figure 3M)**. Conversely, the *s_L_* was 1.4-fold higher for Sox2a **(Figure 3N)**, resulting in *D_1D,Sox2_*=2.63±0.40×10^5^ bp^2^/s and *D_1D,Sox2a_*=5.03±0.86×10^5^ bp^2^/s for Sox2a **(Figure 3O)**, in line with *D_1D_* values determined for LacI^17^.

Together, this indicates that DFR_Sox2_ facilitates target recognition of Sox2 during 1D sliding, whereas DFR_Sox17_ leads to more frequent bypassing of the target site. Sox2 slides more slowly than Sox2a but recognizes its target more effectively, possibly due to a trade-off between scanning speed and recognition efficiency^63,64^. This reveals that DFR_Sox2_ has a strong influence on the ability of Sox2 to recognize the target site during 1D sliding, leading to a more efficient search process compared to DFR_Sox17_.

### Sox TFs interact specifically and nonspecifically with nucleosomes

We next dissected the impact of DFR_Sox2_ on TF-DNA interactions in the context of chromatin. We first examined the binding dynamics of Sox2 and Sox2a to single nucleosomes **(Figure 4A)**. We reconstituted three different mononucleosomes (MNs), using a 250 bp DNA sequence based on the 601WS and recombinant human histone octamers, resulting in MNs flanked by 50 bp linker DNA. One MN contained a Sox motif positioned at superhelical location −6 (SHL-6), a position close to the nucleosome entry site (MN-6), where Sox2 has been found bound with Oct4^65^. A second MN had the motif positioned at SHL+2 (MN+2), an internal position where Sox2 had been shown to bind and locally distort DNA^66^ **(Figure 4B**; see **Methods)**. Finally, we also reconstituted a MN devoid of any binding motif (MN-). We then recorded movies of Sox TFs interacting with these MNs using smTIRF microscopy and constructed 1-CDFs from the resulting fluorescence time-traces **(Figures 4C; S4A-C).**

**Figure 4.**
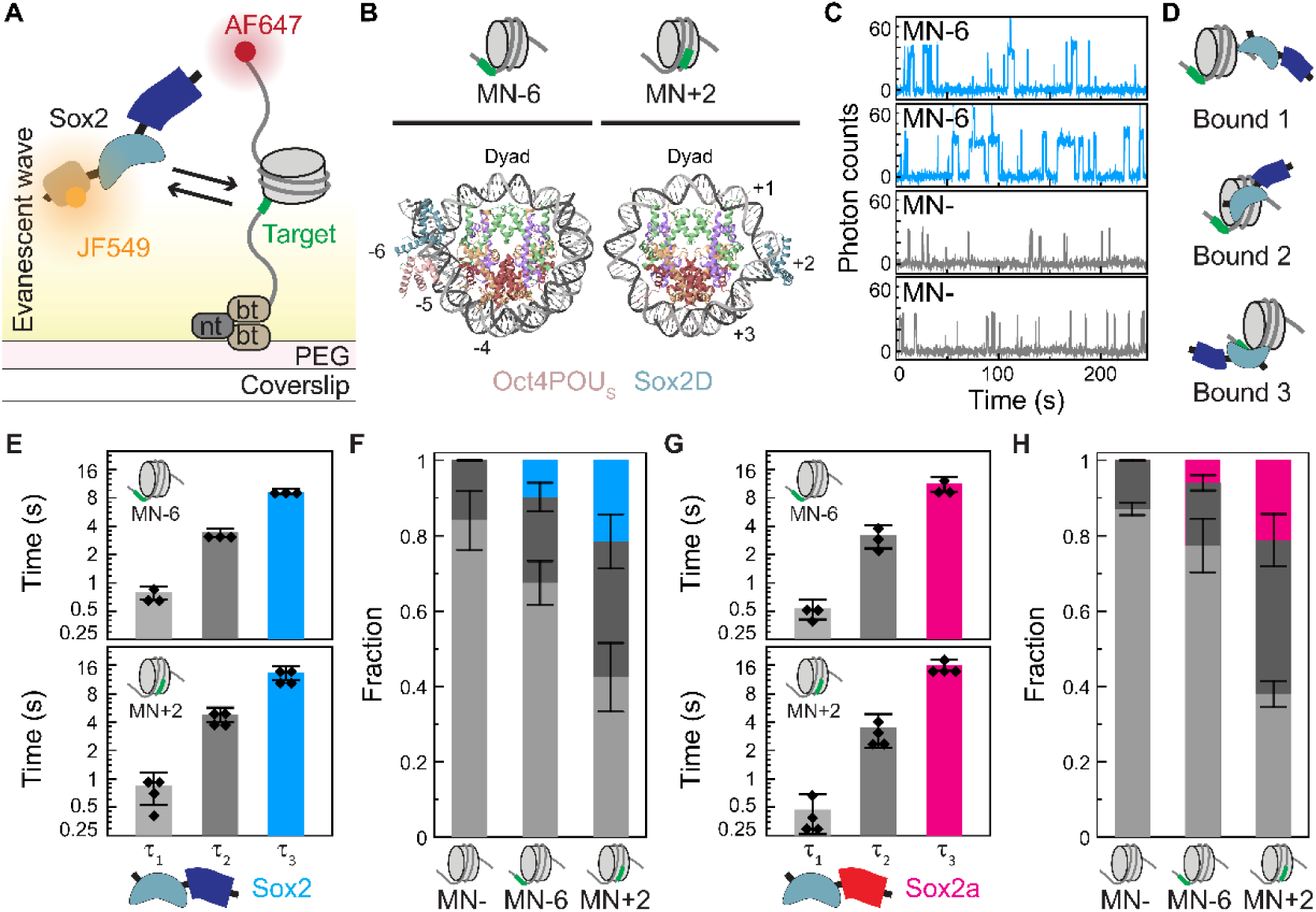
Sox TFs interact specifically and nonspecifically with nucleosomes. **(A)** Scheme of single-molecule experiment to detect Sox TF binding to MNs. **(B)** Structure of nucleosomes bound by Sox2 DBD (Sox2D) together with Oct4 DBD (Oct4POU_s_) at the heterodimer motif located at SHL-6 (left; PDB: 6T90)^65^ and by Sox2D at its motif positioned at SHL+2 (right; PDB: 6T90)_66_. Number: SHL. **(C)** Representative fluorescence time-traces of Sox2 interacting with MN-6 (blue) and MN- (gray). **(D)** Scheme of Sox nucleosome binding modes. States 1 and 2: nonspecific binding to linker DNA and to histones. State 3: Specific binding to target (green). **(E-H)** Residence times (*τ_1_*, gray; *τ_2_*, dark gray; *τ_3_*, blue) and corresponding bound fractions of Sox2 **(E-F)** and Sox2a **(G-H)** on MNs. Symbols: replicates (N=3-4). Bar: mean. Cap: SD.

We found that a triexponential function, associated with three residence times *τ*_1_, *τ*_2_, and *τ*_3_, was required to adequately describe the binding kinetics of both Sox2 and Sox2a on motif-containing MNs **(Figure S4D)**. In contrast, binding kinetics to MN- were well described by a biexponential decay with two residence times *τ*_1_ and *τ*_2_ **(Figure S4D)**. The shortest residence time (*τ*_1,Sox2_=0.8±0.1 s) was observed across all MNs, and being similar to *τ*_1_ in naked DNA, could be attributed to nonspecific DNA interactions **(Figure 4D)**. In contrast, the intermediate residence time (*τ*_2,Sox2,MN-6_=3.4±0.4 s) was not previously seen in free DNA and is thus specific for MNs. As it appeared in both motif-containing and motif-absent MNs, we attributed it to TF-histone interactions **(Figure 4D)**. Finally, the longest residence time (*τ*_3,Sox2,MN-6_=9.3±0.6 s) was unique to motif-containing MNs, confirming its correspondence to specific interactions **(Figure 4D)**.

We then compared Sox2 binding to the motifs within MNs at SHL+2 and SHL-6. While the residence times were comparable for the two positions **(Figure 4E)**, the fraction of specific binding events was higher in MN+2 compared to MN-6, indicating that Sox2 has a higher probability of binding to the internal position at SHL+2 **(Figure 4F)**. This can be explained by the rotational positioning of the motif, which faces outward in MN+2 but inward in MN-6, consistent with previous reports^60^. Finally, a comparison between Sox2 and Sox2a **(Figures 4E-H; Table S4)** showed similar residence times across all tested MNs.

### DFR_Sox2_ increases the fraction of specifically-bound molecules on chromatin

In the cell, pTFs are able to invade compact chromatin^26^. As Sox2 has pioneer activity *in vivo*, we used single-molecule imaging to assess if the nature of the DFR is critical for its chromatin invasion ability^24^ **(Figure 5A)**. We assembled chromatin fibers on DNA templates containing 10 tandem repeats of the 601WS separated by 30 bp linker DNA, either containing a Sox motif at the entry site (SHL-6) of the fifth nucleosome (CT+) or in the absence of any target motif (CT-) **(Figures S5A-C;** see **Methods).**

**Figure 5.**
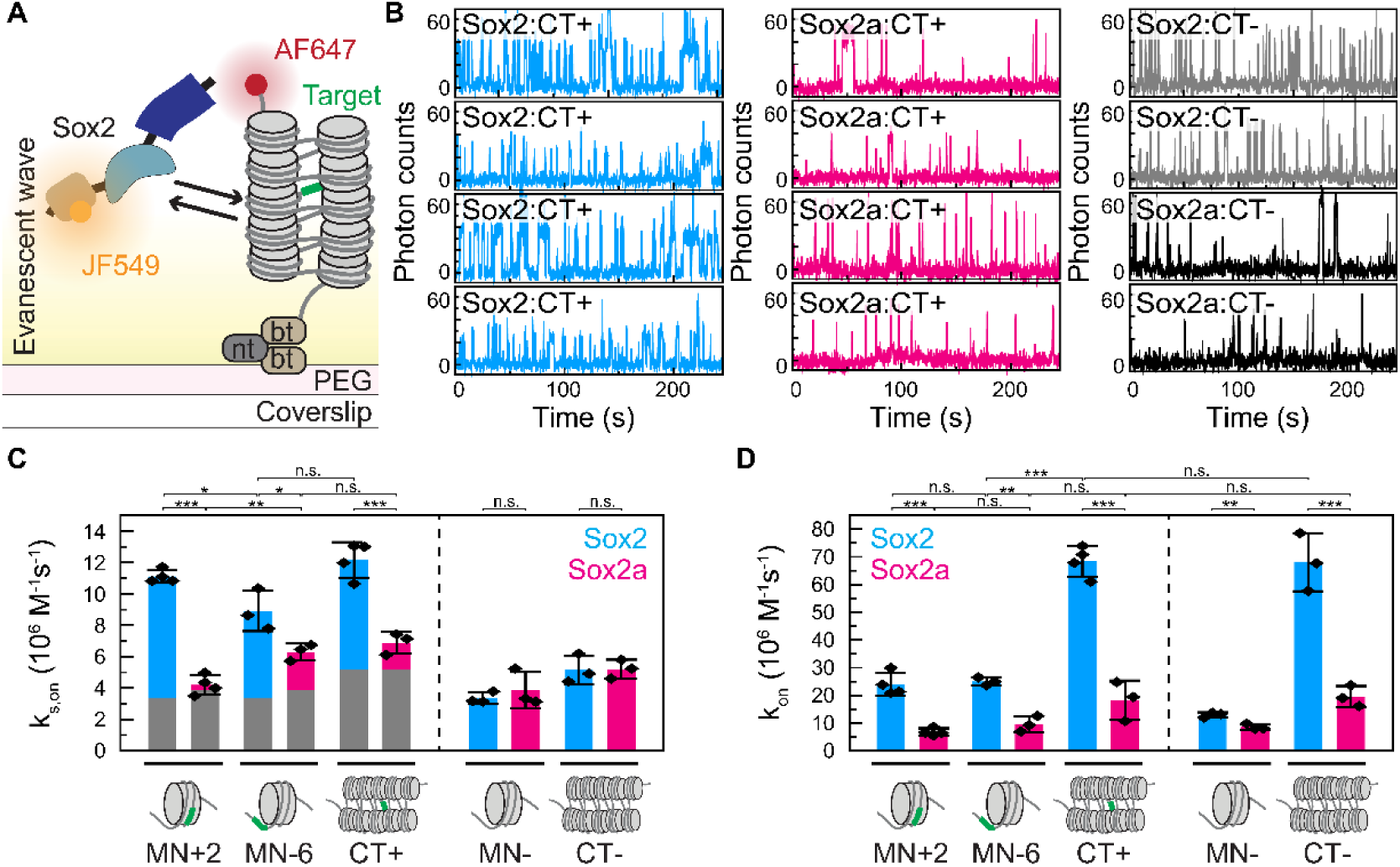
DFR_Sox2_ enhances chromatin invasion through nonspecific interactions with DNA. **(A)** Scheme of single-molecule experiment to detect Sox TF binding to chromatin fibers. **(B)** Fluorescence time-traces of Sox TFs interacting with CT+ and CT-. **(C)** Specific on-rate of Sox TFs to MNs and chromatin (blue: Sox2, red: Sox2a). The background on-rates (gray) measured with control constructs (MN-, CT-) are overlaid with corresponding constructs containing the target site (green). Symbols: replicates (N=3-4). Bar: mean. Cap: SD. **(D)** On-rate of Sox TFs to MNs and chromatin fibers (blue: Sox2, red: Sox2a). Symbols: replicates (N=3-4). Bar: mean. Cap: SD. Statistical significance was assessed using two-tailed t-tests (*: p≤0.05; **: p≤0.01; ***: p≤0.001).

When imaging binding dynamics to these chromatin fibers, Sox2 exhibited more frequent long binding events compared to Sox2a **(Figure 5B)**. Similar to the results obtained with MNs, both Sox TFs showed three different residence times on CT+, while they displayed only two nonspecific binding populations on CT-**(Figures S5D-H)**. The residence times were closely aligned with the identified ranges from the MN dataset **(Table S4)**. On chromatin fibers, nonspecific interactions predominate because of their large size and the consequent challenge for a TF to locate its unique, embedded target site. However, Sox2 exhibited significantly more long binding events (7.0±1.6 %) **(Figure S5F)** compared to Sox2a (1.2±0.3 %) **(Figure S5H)**. This suggests that DFR_Sox2_ enhances Sox2’s ability to access and recognize motifs within chromatin.

The single-molecule binding datasets allowed us to quantify target search efficiency of Sox TFs on MNs and chromatin. To determine *k_s,on_* for Sox2 and Sox2a, we used a residence time threshold (3.8 s for MNs, 3.4 s for chromatin fibers) to select specific binding events, and generate 1-CDF curves of the interspaced search times **(Figures S5I-J)**. We found *k_s,on_* to be markedly higher for Sox2 than for Sox2a on all substrates **(Figure 5C)**. This was especially pronounced for MN+2 compared to MN-6, highlighting Sox2’s superior ability to interact with internal target sites. Of note, we also detected a lower frequency of binding events with residence times above the threshold in the absence of the motif, which originate from the tail of the *τ*_2_ lifetime distribution and could be assigned as a ‘background’ on-rate (shown in grey in **Figure 5C)**.

The CT+ construct contains a central nucleosome with the Sox motif positioned at SHL-6 and further flanked by four and five nucleosomes on either side. In general, a target site within chromatin is expected to be less accessible than when it is located on a nucleosome because of nucleosomes stacking in a tetranucleosome conformation^67–69^. However, the effective *k_s,on_* of Sox2 increased from MN-6 to CT+ by 1.3-fold **(Figure 5C)**, suggesting that 1D diffusion along the chromatin fiber, or a higher local TF concentration due to the availability of many nonspecific binding sites might facilitate target search. In contrast, no such increase in the effective rate was detected for Sox2a. Finally, when including nonspecific binding events **(Figures S5K-L)**, Sox2 exhibited approximately three times higher *k_on_* with all the nucleosome constructs than Sox2a **(Figure 5D)**. As the nature of the DFR did not impact naked DNA on-rate **(Figures S3F-H)**, this suggests that the nonspecific interactions of DFR_Sox2_ increase the on-rate specifically on MNs.

In summary, our on-rate results demonstrate that DFR_Sox2_ promotes prevalent nonspecific binding from 3D diffusion to nucleosomes, enabling Sox2 to invade chromatin more effectively. This ultimately facilitates its target search within chromatin.

### DFR_Sox2_ enhances *in vivo* pioneer activity by facilitating closed chromatin binding and initiation of chromatin opening

We next asked how the different DFRs control chromatin invasion of Sox TFs in cells. We induced Sox2 and Sox2a expression in engineered mESC lines, followed by fluorescence-activated cell sorting (FACS) to ensure similar TF expression levels **(Figure 6A**, **Methods)**. We validated the genome-wide binding profiles of expressed Sox2 and Sox2a against endogenously-expressed Sox2 via Chip-Seq^70^ **(Figures S6A-B)**.

**Figure 6.**
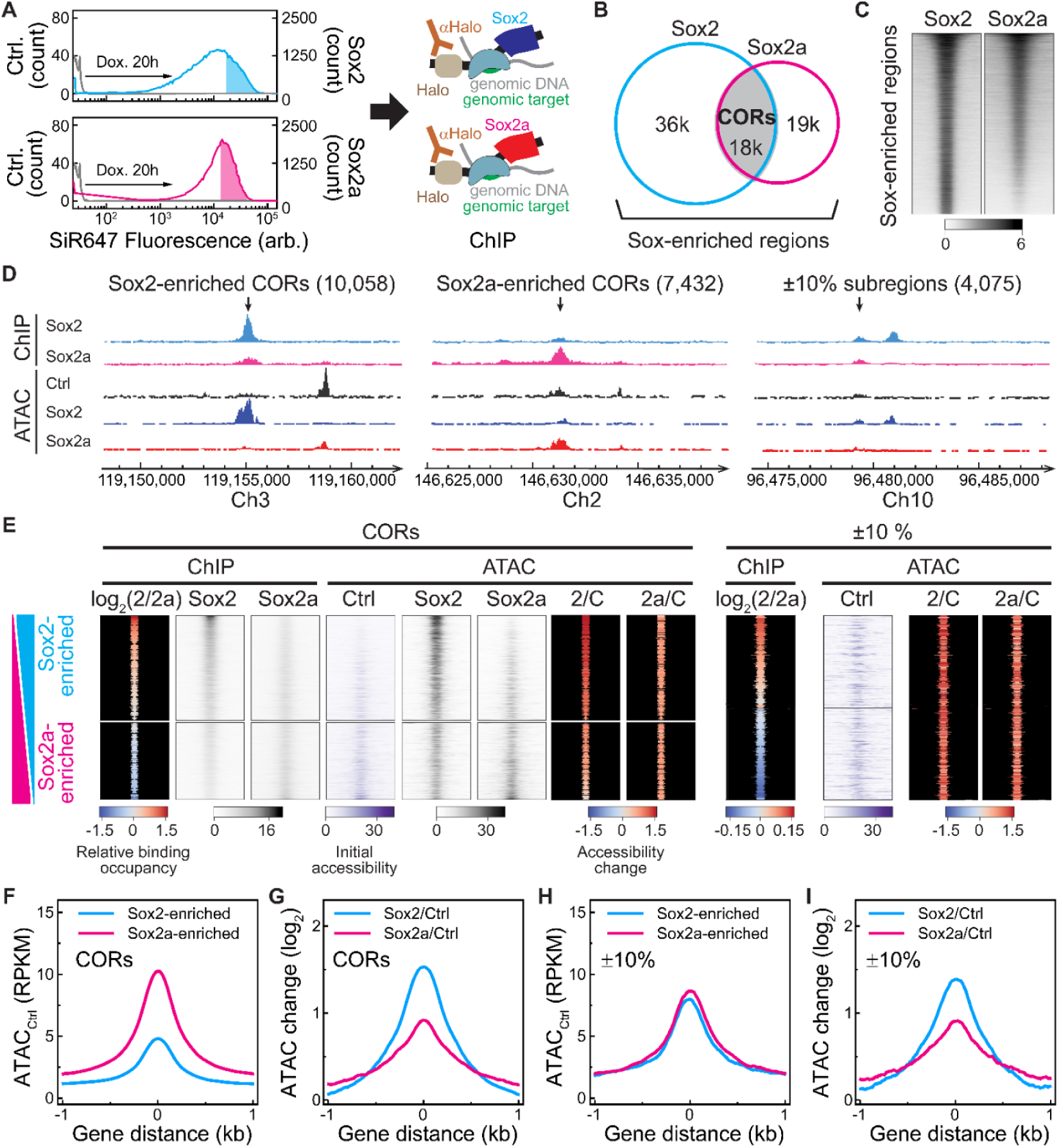
DFR_Sox2_ enhances pioneering activity. **(A)** FACS profiles of mESCs expressing Halo-Sox2 or Halo-Sox2a. Colored areas: matched mean Halo-SiR647 fluorescence intensity. **(B)** Genomic regions enriched by Sox2 (blue) and Sox2a (pink). Two replicates for each Sox TF were merged. **(C)** Heatmaps of ChIP-seq data (RPKM) across Sox-enriched regions. **(D)** Genome track of a genomic region displaying ChIP-seq and ATAC-seq (RPKM) signals, representing Sox2/Sox2a-enriched CORs, and ±10 *%* subregions. The y-axis of Ctrl ATAC is zoomed in threefold compared to the others. The values indicate the number of peaks in each region. **(E)** Heatmaps of ChIP-seq and ATAC-seq data (RPKM) aligned to the log2-fold change in ChIP scores of Sox2 over Sox2a across CORs and ±10 *%* subregions. 2/C: log2-fold change in ATAC scores of Sox2 over Ctrl; 2a/C: log2-fold change in ATAC scores of Sox2a over Ctrl. Log2-fold changes were calculated based on peak signal in bed files. **(F)** Mean score profile of Ctrl ATAC data for Sox2-enriched and Sox2a-enriched CORs. **(G)** Mean score profile of log2-fold change in ATAC scores of Sox TFs over Ctrl across CORs. **(H)** Mean score profile of Ctrl ATAC data for mESCs overexpressing Sox TFs across ±10 *%* subregions. **(I)** Mean score profile of the log2-fold change in ATAC scores of mESCs overexpressing Sox TFs compared to Ctrl mESCs across ±10 *%* subregions.

To quantitatively compare the genomic occupancy of Sox2 and Sox2a, we applied a Drosophila spike-in normalization approach (see **Methods)**. We merged regions enriched for one or both Sox TFs (Sox-enriched regions) **(Figure 6B)**. Sox2a enrichment was well-correlated to Sox2 ChIP scores across Sox-enriched regions **(Figure 6C)**. However, the occupancy level in Reads Per Kilobase Million (RPKM) was higher for Sox2 **(Figure S6C)**. We thus conclude that Sox2a exhibits binding specificity comparable to that of Sox2, despite its lower binding occupancy. When evaluating the overlap between Sox2 and Sox2a ChIP-seq peaks **(Figure 6B)**, we found that a significant proportion of Sox2a-enriched regions did not overlap with those enriched for Sox2, and the overall number of occupied regions was lower for Sox2a. Altogether, these results are in line with the higher specific on-rate of Sox2 compared to Sox2a previously observed in both *in vivo* and *in vitro* SMT analysis, and directly show that the DFR controls target site selection in cells.

To analyze genome-wide binding patterns, we split Sox-enriched regions into two groups **(Figure 6B)**: i) commonly occupied regions (CORs) containing ChIP-seq peaks for both Sox2 and Sox2a, and ii) exclusive regions where either Sox TF displayed ChIP-seq peaks. We then plotted the log2-fold change of the COR ChIP scores for Sox2 over Sox2a in descending order. CORs were further sub-clustered into three groups defined by i) higher Sox2 binding; ii) higher Sox2a binding; iii) equal binding of Sox2 and Sox2a within a ±10% margin **(Figure 6D)**. The larger number of Sox2-enriched CORs **(Figure 6D)** and higher ChIP enrichment of Sox2 across CORs **(Figure S6D)** confirmed that Sox2 exhibited greater genomic occupancy compared to Sox2a. Besides, motif analysis across exclusive regions and CORs revealed comparable binding specificity of Sox2 and Sox2a **(Figures S6E-F)**.

To assess the ability of Sox2 and Sox2a to open compact regions upon binding, we generated ATAC-seq data for the same cell lines. We aligned ATAC-seq profiles with ChIP log2-fold changes to assess chromatin accessibility across CORs. CGR8 cells expressing only H2B-SNAP (Ctrl) served as a control to measure initial chromatin accessibility **(Figure 6E)**. mESCs overexpressing Sox2 exhibited higher chromatin accessibility at CORs than those overexpressing Sox2a **(Figure S6G)**. However, Ctrl ATAC-seq revealed that Sox2-enriched CORs were initially 1.7-fold more closed compared to Sox2a-enriched CORs **(Figure 6F)**. This confirms that DFR_Sox2_ enhances chromatin invasion of Sox2, as we observed *in vitro*. We then computed the log2-fold change of ATAC scores for Sox2 and Sox2a over Ctrl. Sox2 increased chromatin accessibility across CORs more than Sox2a **(Figures 6G, S6H)**.

Finally, we determined changes in chromatin accessibility in regions with comparable Sox2 and Sox2a occupancy (within a range of ±10 %) and initial chromatin accessibility **(Figure 6H)**. Across these regions, chromatin accessibility increased more for Sox2 than for Sox2a **(Figure 6I)**. Altogether, this data indicates that DFR_Sox2_ allows Sox2 to bind to poorly accessible chromatin and to open bound regions more effectively, which endows Sox2 with superior pioneering activity.

## DISCUSSION

TF target search has been described using the facilitated diffusion model, which combines 3D diffusion and 1D sliding on DNA to locate specific binding sites^11,12,71,72^. Bacterial TFs, such as LacI in E. coli, are reliant on 1D diffusion^15,62^. In contrast, eukaryotic TFs search the genome within a highly complex nuclear environment, characterized by chromatinized DNA organized into higher-order structures including chromatin fibers, loops^73,74^, TADs and A/B compartments^75^. Pioneer TFs such as Sox2 evolved mechanisms to penetrate compact chromatin states, to search the underlying DNA and to open chromatin structure after target motif recognition^25,36,53^. Earlier *in vivo* single-molecule imaging revealed that Sox2 spends significant time in free 3D diffusion^2^, which may reflect a generic strategy used by eukaryotic TFs to overcome obstacles such as nucleosomes that prevent sliding^76^.

A major challenge in understanding gene regulation is uncovering how the biochemical properties of TFs shape target search efficiency, particularly given the role of large disordered regions in search dynamics, target selection, and overall TF function^77^. Although these domains lack a defined structure, both short sequence elements and broader physicochemical properties (e.g., charge and hydrophobicity), particularly in the DFR, modulate the DNA− binding characteristics of the TF DBD. These have significant implications, as studies in yeast^27,78,79^ and mammalian cells^32,35,53,80,81^ demonstrate that the DBD alone is insufficient for target selection, and that truncating the DFR alters site specificity. Moreover, diffusional properties in the nucleus are modified upon swapping or deleting DFRs, including a loss of diffusional confinement^53,82^ or scanning of compact chromatin regions^35,78^. To understand how the DFR impacts target search, we mechanistically dissected this process for Sox2 within the chromatinized genome, combining *in vitro* and *in cell* biophysics and genomics.

We propose that the electrostatic properties of the DFRs plays a key role in target search. DFR_Sox17_, which is associated with less efficient target search, is more negatively charged and results in weaker DNA retention of TFs, in particular when challenged with high salt concentrations, pointing towards an electrostatic mechanism. These results agree with our previous findings that more positively charged TFs exhibit higher search efficiency^40^. Of note, negatively-charged IDRs were also reported to control nonspecific DNA binding through an autoinhibition mechanism^10,79^, but Sox2 lacks the necessary domain^80^. In addition, we cannot exclude other stabilizing mechanisms related to specific sequence elements within the about 100 amino acid-long DFR. In addition, in all of our experiments we have focused on single TFs interacting with DNA or chromatin, thus we are blind to potential changes in the interactions with other proteins mediated by the DFR. At high local concentrations, Sox2 has the propensity to form condensates^81,83^, which increases local retention and may play an important role in the activation of super-enhancer*s*^84,85^. Although IDRs often contribute to condensate formation, the on-rate enhancing effects of single TFs identified here appear independent of such condensation processes.

We compare the effects of DFRs from Sox2 and Sox17 that show highly differential mitotic chromatin binding^40^, albeit having a homologous DBD recognizing a very similar DNA motif. We revealed distinct mechanisms by which DFR_Sox2_ and DFR_Sox17_ control TF target search on DNA versus chromatin **(Figure 7A)**. On naked DNA, DFR_Sox2_ drives rapid target recognition during 1D sliding, whereas DFR_Sox17_ results in an increased sliding length but more frequent target site bypassing. In contrast, on chromatinized DNA, DFR_Sox2_ promotes frequent nonspecific interactions with both histones and nucleosomal DNA which can transition into specific target binding. Moreover, DFR_Sox2_ facilitates chromatin invasion, by enabling the TF to access DNA sites embedded within compact chromatin **(Figure 7B)**. Conversely, DFR_Sox17_ is associated with a lower overall on-rate and thus fewer nonspecific binding events, resulting in a strongly reduced chromatin binding ability. This dual function of DFR_Sox2_ thereby drives efficient target search within the complex chromatin environment, driving potent pioneer activity of Sox2 for local chromatin accessibility control **(Figure 7B)**.

**Figure 7.**
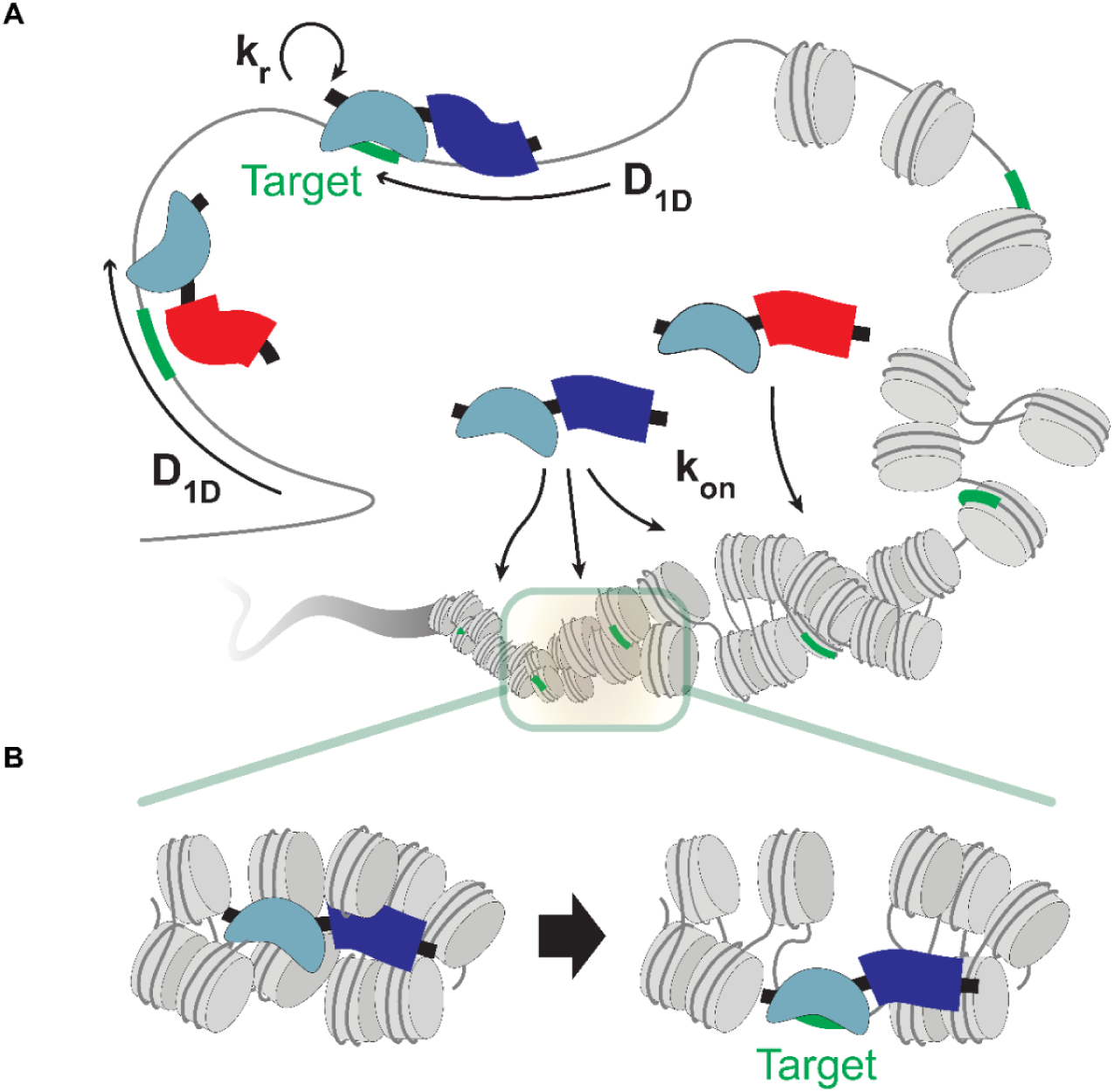
Eukaryotic TF target search model on chromatinized genome via electrostatic interactions. (A) Distinct target search mechanism on chromatinized versus naked DNA. (B) DFR_Sox2_ enhances the abilities of chromatin invasion and local chromatin opening.

In summary, our results show that rapid, transient nonspecific chromatin binding events, relying on a detailed charge balance, allow Sox2 to sample the chromatinized genome efficiently. This may be a common principle extending to a broad range of eukaryotic TFs, in particular pioneer TFs that need to access compact chromatin regions.

## ACKNOWLEDGEMENTS

We thank Cedric Deluz for providing pLV-PGK-H2B-SNAP plasmid. Dr. Luc Reymond for providing the synthesized Halo-SiR647 dye. We are grateful to Dr. Armelle Tollenaere for her assistance with sequencing sample preparation. We also thank Dr. Bastien Mangeat and Elisa Cora at the Gene Expression Core Facility at EPFL for performing next-generation sequencing, and Dr. Miguel Garcia, Dr. Valérie Glutz, and Dr. Francesco Palumbo at the Flow Cytometry Core Facility for their support with FACS. Our appreciation extends to Dr. Florence Pojer, Dr. Kelvin Lau, and Dr. Amédé Noredine Larabi at the Protein Production and Structure Core Facility at EPFL for their help with baculovirus generation and expression of Sox2 and Sox2a constructs. We acknowledge our colleagues in LCBM for expressing the histone octamers, and we thank the Bioimaging and Optics Core Facility at EPFL for maintaining the Zeiss LSM700 UP2 and Nikon-CSU-W1 microscopes. This work was supported by the Swiss National Science Foundation (310030_200604 to B.F.), (EPFL annual dotation to D.S.), as well as the interdisciplinary PhD program from the School of Life Sciences, EPFL.

## AUTHORS CONTRIBUTIONS

Conceptualization: BF, DS and SS; Methodology: SS; Formal Analysis: SS; Investigation SS; Visualization: SS; Resources: SS; Supervision: BF and DS; Writing: BF, DS and SS; Funding acquisition: BF and DS.

## DECLARATION OF INTERESTS

The authors declare no competing interests.

## SUPPLEMENTARY INFORMATION

Figures S1 to S6 Tables S1 to S4

Tables S5 to S12, which include detailed experimental and data processing settings as well as material information, are provided in **STAR★ METHODS**.

## STAR★ METHODS

### KEY RESOURCES TABLE

| Antibodies, Chemicals, and Recombinant Proteins |  |  |
| --- | --- | --- |
| Reagent/Resource | Source | RRID |
| Halo-Trap Agarose beads | ChromoTek, ota |  |
| Rabbit anti-NANOG antibody | CST, 8822 |  |
| Chicken anti-Rabbit IgG antibody | Invitrogen, A21443 |  |
| JaneliaFluor-549 Halo | Promega, GA1110 |  |
| AlexaFluor-647 NHSEster | Invitrogen, A20006 |  |
| XFD647 NHSEster | AATbio, 1833 |  |
| T4 DNA Ligase | NEB, M0202 |  |
| BglII | NEB, R0144 |  |
| BamHI-HF | NEB, R3136 |  |
| EcoRI-HF | NEB, R3101 |  |
| DraIII-HF | NEB, R3510 |  |
| EcoRV-HF | NEB, R3195 |  |
| BsaI-HFv2 | NEB, R3733 |  |
| ScaI-HF | NEB, R3122 |  |
| Quick CIP | NEB, M0525 |  |
| Cell lines |  |  |
| Reagent/Resource | Source | RRID |
| CGR8 (ESC Mus musculus) | Sigma-Aldrich, 07032901 | N/A |
| 2TS22C (ESC Mus musculus) | Masui et al., 2007 | N/A |
| HEK293T | ATCC | CVCL_0063 |
| SES4.1 (ESC Mus musculus) | This study | N/A |
| SES5.1 (ESC Mus musculus) | This study | N/A |
| SES6.1 (ESC Mus musculus) | This study | N/A |
| SES7.1 (ESC Mus musculus) | This study | N/A |
| SES8.1 (ESC Mus musculus) | This study | N/A |
| SES9.1 (ESC Mus musculus) | This study | N/A |
| SES11 (ESC Mus musculus) | This study | N/A |
| SES13 (ESC Mus musculus) | This study | N/A |
| SES14 (ESC Mus musculus) | This study | N/A |
| DH5a (E. coli) | NEB, C2987H | N/A |
| Hi5 | PTPSP, EPFL | N/A |
| Sf9 | PTPSP, EPFL | N/A |
| Plasmid DNA |  |  |
| Reagent/Resource | Source | RRID |
| psPAX2 | Addgene | Addgene_12260 |
| pMD2.G | Addgene | Addgene_12259 |
| pLV-TRE3G-Halo-Sox2-Puro | Deluz et al., 2016 | N/A |
| pLV-PGK-rtTA3G-Bsd | Deluz et al., 2016 | N/A |
| recP1(pWM531-EcoRI-DraIII-1×601WS-BamHI-4×601WS-BglII-BsaI) | Kilic et al., 2018 | N/A |
| Spla5 (pLV-TRE3G-Halo-Sox2a-Puro) | This study | N/A |
| Spla7 (pLV-TRE3G-Halo-Sox17-Puro) | This study | N/A |

| Spla9 (pLV-TRE3G-Halo-Sox17b-Puro) | This study | N/A |
| --- | --- | --- |
| Spla11 (pLV-PGK-H2B-SNAP) | This study | N/A |
| Spla14 (PaceBac1-Hisx8-Halo-Sox2-TwinStrep) | This study | N/A |
| Spla16 (PaceBac1-Hisx8-Halo-Sox2-TwinStrep) | This study | N/A |
| Spla22 (pLV-TRE3G-Halo-Sox2D-Puro) | This study | N/A |
| Spla23 (pLV-TRE3G-Halo-Sox17D-Puro) | This study | N/A |
| Spla30 (pLV-EF1a-Halo-Sox2-Puro) | This study | N/A |
| Spla31 (pLV-EF1a-Halo-Sox2a-Puro) | This study | N/A |
| Spla35 (pWM531-EcoRI-10×601WS-BsaI) | This study | N/A |
| Spla36 (pMARQ-BglII-601WS-DraIII) | This study | N/A |
| Spla37 (pMARQ-BglII-Sox601WS-DraIII) | This study | N/A |
| Spla38 (pWM531-EcoRI-4×601WS-Sox601WS-4×601WS-BsaI) | This study | N/A |
| Spla42 (pMARQ-EcoRI-4×601WS-Sox601WS-DraIII) | This study | N/A |
| Spla43 (pMARQ-EcoRI-5×601WS-DraIII) | This study | N/A |
| **double-stranded DNA** |  |  |
| Reagent/Resource | Source | RRID |
| Sdna3 (50bp naked DNA containing a Sox motif) | This study | N/A |
| Sdna4 (100bp naked DNA containing a Sox motif) | This study | N/A |
| Sdna5 (50bp naked DNA based on 601WS sequence) | This study | N/A |
| Sdna6 (30bp naked DNA containing a Sox motif) | This study | N/A |
| Sdna8 (25bp naked DNA containing a Sox motif) | This study | N/A |
| Sdna9 (250bp naked DNA containing a Sox motif) | This study | N/A |
| Sdna10 (250 bp; MN-6, nucleosomal DNA) | This study | N/A |
| Sdna11 (250 bp; MN+2, nucleosomal DNA) | This study | N/A |
| Sdna16 (250 bp; MN-, nucleosomal DNA) | This study | N/A |
| Sdna17 (CT+, chromatin DNA) | This study | N/A |
| Sdna18 (CT-, chromatin DNA) | This study | N/A |
| Sdna19 (MMTV buffer DNA) | This study | N/A |
| **Star oligonucleotides** |  |  |
| Reagent/Resource | Source | RRID |
| Soli175 (Sanc1 forward) | This study | N/A |
| Soli176 (Sanc1 reverse) | This study | N/A |
| Soli179 (reverse; 5'-BamHI-EcoRI-3') | This study | N/A |
| **Software and algorithms** |  |  |
| Reagent/Resource | Source | RRID |
| MATLAB R2023b | Mathworks | SCR_001622 |
| Custom MATLAB scripts | This study | N/A |
| Origin 2022 | OriginLab | SCR_014212 |
| FiJi Version 2.14.0/1.54f | Schindelin et al., 2012 | SCR_002285 |
| TrackIt Version 1.4 | Khun et al., 2021 | N/A |
| Spot-On Version MATLAB | Hansel et al., 2018 | N/A |
| StarDist Fiji plugin | Schmidt et al., 2018 | N/A |
| STAR Version 2.7.6a | Dobin et al., 2013 | SCR_015899 |
| Picard Version 2.23.7 | Broad Institute | SCR_006525 |
| MACS Version 3.0.0a7 | Zhang et al., 2008 | SCR_013291 |
| DeepTools Version 3.3.0 | Ramírez et al., 2016 | SCR_016366 |
| BEDtools Version 2.30.0 | Quinlan and Hall, 2010 | SCR_006646 |
| SAMtools Version 1.14 | Li et al., 2009 | SCR_002105 |
| HOMER2 Version 4.11 | Heinz et al., 2010 | SCR_010881 |
| R Version 4.1.2 | R Project | SCR_001905 |
| R studio Version 2021.09.2+382 | Rstudio | SCR_000432 |
| NIS-Elements AR | Nikon | SCR_014329 |
| Image Lab Version 4.1 | Bio-rad | SCR_014210 |

### LEAD CONTACTS AND MATERIALS AVAILABILITY

Plasmids and cell lines are available upon request to the Lead Contacts. Further information and requests for resources and reagents should be directed to and will be fulfilled by the Lead Contacts, Beat Fierz or David Suter.

### EXPERIMENTAL MODEL AND SUBJECT DETAILS

#### mESC culture

mESCs were routinely cultured on 0.1% gelatin-coated (Sigma, G9391) 100 mm Petri dishes at 37 °C with 5% CO2 in GMEM (Sigma, G5154-500ML), supplemented with 10% ESC-qualified fetal bovine serum (Gibco, 16141-079), 1% nonessential amino acids (Gibco, 11140-050), 2 mM L-glutamine (Gibco, 25030-024), 2 mM sodium pyruvate (Sigma, S8636-100ML), 100 μM 2-mercaptoethanol (Sigma, 63689-25), 1% penicillin-streptomycin (BioConcept, 4-01 F00-H), in-house produced leukemia inhibitory factor (LIF), CHIR99021 (Merck, 361559-5MG) at 3 μM and PD184352 (Sigma, PZ0181-25MG) at 0.8 μM. Cells were passaged every two to three days using trypsinization (Sigma, T4049-100ML). Once cells reached 60-70% confluency, one-sixth to one-tenth of the culture was used for the next passage.

For imaging, one day prior to measurement, 170 μm glass-bottom 35 mm Petri dishes were pre-coated with a 1:10 dilution of Biolaminin (BioLamina, LN511-0202) in DPBS containing magnesium and calcium ions (Gibco, 14040117). Cells were then plated on the dishes in FluoroBrite DMEM (ThermoFisher, A18967-01) supplemented with 10% ESC-qualified fetal bovine serum, 1% non-essential amino acids, 2 mM L-glutamine, 2 mM sodium pyruvate, 100 µM 2-mercaptoethanol (Sigma-Aldrich, 63689-25ML-F) 1% penicillin/streptomycin, LIF, CHIR99021 at 3 µM and PD184352 at 0.8 µM.

#### Lentiviral vector production

Lentiviral vectors were produced by transfection of HEK 293T cells with the envelope (psPAX2, Addgene, 12260), packaging (pMD2.G, Addgene 12259), and lentiviral construct of interest using calcium phosphate transfection^86,87^. The HEK293T cells were cultured in GlutaMAX-containing DMEM (Gibco, 31966021) with 10% fetal bovine serum (Gibco, 10270106) and 1% penicillin-streptomycin (BioConcept, 4-01F00-H). After two days post-transfection when they reach 80% confluency, the virus-containing medium was harvested and cleared of cell debris by ultracentrifugation. Viral vectors were concentrated 120-fold by ultracentrifugation at 20,000 x g for 120 min at 4°C. 50,000 cells in 1 mL of medium in a 12-well plate were transduced with 50 μl of concentrated lentiviral vector particles to generate stable cell lines.

#### Generation of dox-inducible Halo-Sox TF expressing mESCs

**Table S5.** Halo-Sox TF expressing mESCs.

| mESCs | pLV-TRE3G-Halo-Sox | pLV-PGK-rtTA3G | pLV-pGK-H2B-SNAP |
| --- | --- | --- | --- |
| **SES4.1** | Sox2 | ○ | ○ |
| **SES5.1** | Sox2a | ○ | ○ |
| **SES6.1** | Sox2D | ○ | ○ |
| **SES7.1** | Sox17 | ○ | ○ |
| **SES8.1** | Sox17D | ○ | ○ |
| **SES9.1** | Sox17b | ○ | ○ |
| **SES11 (Ctrl.)** | - | - | ○ |

Sox TF-expressing embryonic stem (SES) cell lines in Table S5 are mESC derivatives of CGR8 cells engineered to express Halo-Sox TFs upon doxycycline induction. Three transgenes were inserted into the genome using a lentiviral vector approach. The lentiviral plasmids (pLV-TRE3G-Halo-Sox) were generated using the pLV-TRE3G-Halo-Sox2 plasmid^88^ and cDNA sequence of Sox17^40^, which were sourced from the laboratory. Using the In-fusion cloning technique (Takara Bio, 638947), we generated lentiviral plasmids containing transgenes for every Halo-Sox TF. The obtained products were transformed into HB101 competent cells. Clones were sequenced to verify the presence of the sequence of interest.

The genes encoding the recombinant proteins, Halo-Sox TFs (sequences in Table S6), regulated by seven Tet operators, and H2B-SNAP driven by an hPGK promoter, were integrated into the mouse genome of SES cells as indicated in Table S5. CGR8 cells (Sigma, 07032901-1VL) were transduced to constitutively express rtTA3G, Halo-Sox TFs, or H2B-SNAP according to Table S7. To select for SES cells containing the transgene of interest, SES cells were treated with 2 µg/mL of puromycin (Gibco, A11138-03) for TRE3G-Halo-Sox2 transgene and/or 1 µg/mL of blasticidin for PGK-rtTA3G, for a week. PGK-H2B-SNAP positive SES cells were sorted by FACS thresholding SNAP-SiR647 intensity after 30 minute-incubation of SNAP-labeling SiR647 dye (NEB, S9102S) in the cells.

**Table S6.**
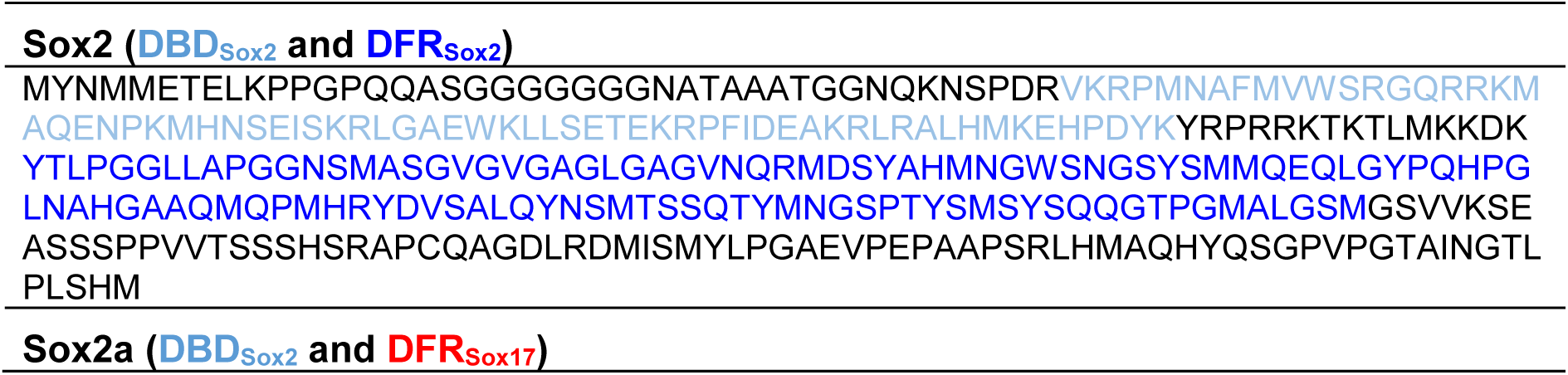
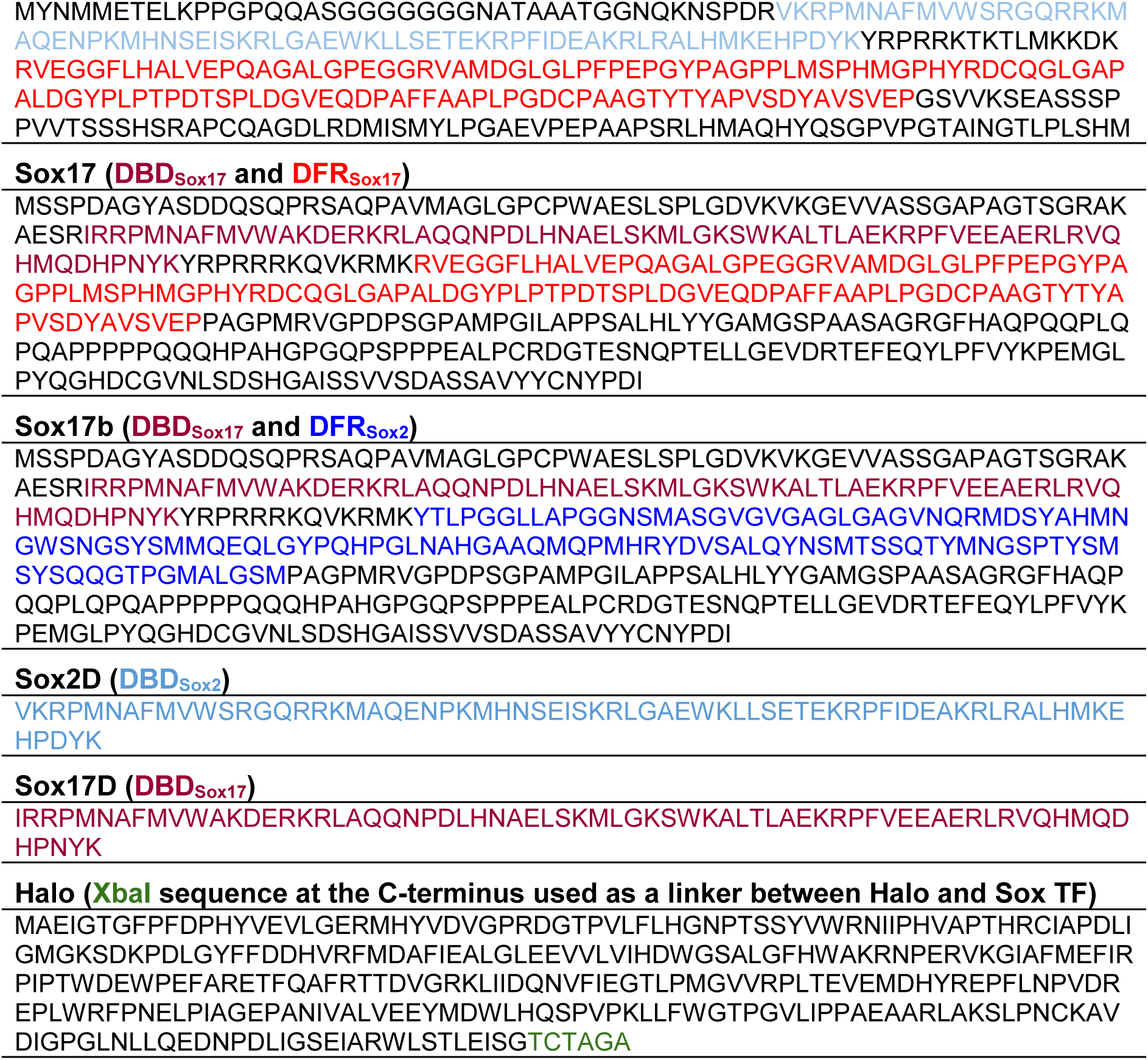
amino acid sequences of Sox TFs and Halo-tag.

#### Generation of 2TS22C cells expressing Halo-Sox2 or Halo-Sox2a

The 2TS22C cell line^42^ enables knockdown of endogenous Sox2 upon dox treatment. Similar to the dox-inducible Halo-Sox expressing mESCs (Table S5), 2TS22C cells expressing Halo-Sox2 (SES13) or Halo-Sox2a (SES14) were generated using lentiviral vector transduction. To generate pLV-EF1α-Halo-Sox plasmids, mCherry in the pLV-EF1α-mCherry plasmid fragment was exchanged with Halo-Sox2 or Halo-Sox2a using restriction cloning. Finally, SES13 and SES14 were sorted by FACS based on Halo-SiR647 intensity after a 30 minute-incubation with 50 nM of Halo-SiR647 dye at 37 °C.

### METHOD DETAILS

#### Pluripotency maintenance assay

To ensure similar expression levels of Halo-Sox2 and Halo-Sox2a, SES13 and SES14 were incubated with Halo-labeling SiR647 dye for 30 minutes at 37 °C following by FACS using the intensity gate, resulting in the same mean fluorescence intensity between SES13 and SES14 samples. Immediately after FACS, 1000 cells were seeded on gelatin-coated, 170 μm glass-bottom 35 mm Petri dishes in GMEM (Sigma, G5154-500ML), supplemented with 10% ESC-qualified fetal bovine serum (Gibco, 16141-079), 1% nonessential amino acids (Gibco, 11140-050), 2 mM L-glutamine (Gibco, 25030-024), 2 mM sodium pyruvate (Sigma, S8636-100ML), 100 µM 2-mercaptoethanol (Sigma, 63689-25), 1% penicillin-streptomycin (BioConcept, 4-01 F00-H), LIF, CHIR99021 (Merck, 361559-5MG) at 3 µM and PD184352 (Sigma, PZ0181-25MG) at 0.8 µM. Following this, 1 µg/mL of dox was added to induce knockdown of endogenous Sox2. The medium supplemented with dox was changed on the third and fifth day after seeding.

On the sixth day after dox addition, cells were fixed with 4% formaldehyde (ThermoFisher, 28908) for 10 minutes at room temperature, gently washed with DPBS (Gibco, 14040091), and stored at 4 °C in 2 mL of DPBS. Since differentiation of mESCs leads to colony flattening and loss or regular edges^89–91^, colony morphology serves as a metric of the pluripotency status. The shape of cell colonies was assessed using phase-contrast microscopy (Zeiss LSM700 UP2, BIOP, EPFL) with a 10x air objective (Objective EC Plan-Neofluar 10x/0.3 Ph1 M27) on an inverted Zeiss Axio Observer Z1 microscope. On the fifth day after dox addition, 170 μm glass-bottom 35 mm Petri dishes were pre-coated with a 1:10 dilution of Biolaminin (BioLamina, LN511-0202) in DPBS containing magnesium and calcium ions (Gibco, 14040117). Then, for NANOG immunofluorescence imaging, 600 cells were seeded on the dishes with fresh medium supplemented with dox. After cell fixation on day 6 using 4% formaldehyde (ThermoFisher, 28908), cells were permeabilized with 0.5% Triton (AppliChem, A1388,0500) at room temperature, followed by multiple washes using DPBS (Gibco, 14040091). Blocking was performed with 1% BSA solution (Sigma-Aldrich, A7906; in DPBS) for 30 minutes. Subsequently, a rabbit anti-NANOG antibody (CST, 8822) was diluted 1:500 in 1% BSA and incubated with the cells overnight at 4 °C. After washing the antibody off with DPBS, cells were incubated for 1 hour at room temperature with AF647-conjugated secondary Chicken anti-Rabbit IgG antibody (Invitrogen, A-21443) at a 1:500 dilution in 1% BSA solution. The cells were washed gently twice with 0.1% Tween-20 (Fisher Scientific, 10113103; in DPBS) and once with DPBS. Finally, Fluoromount G with DAPI (SouthernBiotech, 0100-20) was applied to the samples, and cells were incubated for at least 10 minutes before imaging.

Imaging was performed on Inverted Nikon Eclipse Ti2-E motorized microscope using confocal microscopy (Nikon-CSU-W1, BIOP, EPFL) with a 60x oil objective (Objective CFI Plan Apo Lambda 60x/1.40). 638 nm and 405 nm diode lasers were used to excite AF647 dye and DAPI, respectively. For quantification, the StarDist plugin^92^ in Fiji^93^ was employed to detect nuclei based on DAPI signals. AF647-NANOG IF intensity was then measured within the nuclei of each cell.

#### In vivo SMT sample preparation

**Table S7.** Dox and Halo-labeling JF549 concentrations for in vivo SMT.

|  | Sox2 | Sox2a | Sox17 | Sox17b | Sox2D | Sox17D |
| --- | --- | --- | --- | --- | --- | --- |
|  | **Preparation to measure residence times** |  |  |  |  |  |
| **Dox (ng/ml)** | 5-10 | 5-10 | 25-30 | 25-35 | 20 | 30 |
| **Halo-JF549 (pM)** | 5-10 | 2.5-15 | 2.5-50, 1000 | 10-35 | 20 | 30 |
|  | **Preparation to measure diffusing dynamics** |  |  |  |  |  |
| **Dox (ng/ml)** | 10 | 10 | 25-30 | 25-30 |  |  |
| **Halo-JF549 (pM)** | 10 | 10 | 25-30 | 25-30 |  |  |

One day prior to imaging, Approximately 10^5^ cells were plated on a 170 μm glass-bottom 35 mm Petri dish in FluoroBrite DMEM (ThermoFisher, A18967-01) supplemented with 10% ESC-qualified fetal bovine serum (Gibco, 16141-079), 1% nonessential amino acids (Gibco, 11140-050), 2 mM L-glutamine (Gibco, 25030-024), 2 mM sodium pyruvate (Sigma, S8636-100ML), 100 μM 2-mercaptoethanol (Sigma, 63689-25), 1% penicillin-streptomycin (BioConcept, 4-01 F00-H), LIF, CHIR99021 (Merck, 361559-5MG) at 3 µM and PD184352 (Sigma, PZ0181-25MG) at 0.8 µM. The dishes were already pre-coated with a 1:10 dilution of Biolaminin (BioLamina, LN511-0202) in DPBS containing magnesium and calcium ions (Gibco, 14040117).

The specific concentrations of dyes and dox to visualize Halo-Sox molecules were optimized for each Halo-Sox TF. Cells were incubated overnight with dox to induce Halo-Sox TF expression. Before the measurement, we performed Halo-labeling with the JF549 dye (Promega, GA1110) to measure the residence times and diffusing dynamics. Dox and Halo-JF549 concentrations are indicated in Table S7. For all cell lines and conditions, 1-2 nM of SNAP-labeling SiR647 (NEB, S9102S) were applied. The dyes were freshly incubated for 30 minutes at 37 °C to visualize Halo-Sox TFs and SNAP-H2B, respectively, before imaging. Excess dye was gently washed out using the imaging medium.

#### In vivo smHILO microscopy setup

An inverted Nikon Eclipse Ti2-E fluorescence microscope equipped with a 100× oil immersion NA 1.49 objective (SR HP Apo TIRF) was employed. Fluorophore excitation was achieved using the N-STORM Direct Point illuminator. Two-color imaging was performed with a 30 mW 561 nm diode laser for Halo-JF549 excitation, and a 50 mW 638 nm diode laser for SNAP-SiR647 excitation, with a 405/488/568/647 DM wheel and DM 568 LP emission filter. Dye emission was detected using two Photometrics Prime 95B sCMOS cameras with a physical pixel size of 11×11 mm^2^.

**Table S8.** Number of single-cell movies.

| **Time interval (ms)** | **Sox2** | **Sox2a** | **Sox17** | **Sox17b** | **Sox2D** | **Sox17D** |
| --- | --- | --- | --- | --- | --- | --- |
| **120** | 70 | 87 | 83 | 54 | 85 | 26 |
| **240** | 42 | 51 | 22 | 35 | 80 | 25 |
| **480** | 19 | 41 | 30 | 35 | 56 | 17 |
| **960** | 38 | 44 | 29 | 35 | 60 | 23 |
| **Total** | 169 | 223 | 164 | 159 | 281 | 91 |
| **Continuous (100 Hz)** | 86 | 38 | 38 | 72 | - | - |

Four different time intervals were designed to measure the residence times, varying dark times (120, 240, 480, and 960 ms) while maintaining a 60-ms illumination period. This multi-time interval measurement allowed for the observation of longer binding times (>100 s) and enabled correction for photobleaching, since the time-lapse imaging conditions only affect the photobleaching rate without altering residence times. The number of cells per time interval and Sox TF is indicated in Table S8. SNAP-H2B was imaged with a 60-ms exposure time once every 9–10 frames of Halo-JF549.

To measure diffusion dynamics, we recorded continuous time-lapse movies for 30 seconds with a 100 Hz frame rate (cell numbers are indicated in Table S8). Prior to capturing Sox TF kinetics, a snapshot of SNAP-H2B was taken with a 100 ms exposure. We used the same microscope settings as for diffusion analysis, except for the laser power for JF549 activation that was increased to 110 mW.

#### Expression, labeling, and purification of Sox TFs

Each Halo-Sox TF construct contained a twin-Strep tag at the C-terminus and an 8×His tag at the N-terminus, and was cloned into the donor plasmid pACEBac1 (bearing a gentamicin resistance marker and a YFP reporter gene). Baculovirus production from Sf9 cells and subsequent Halo-Sox2 expression in Hi5 cells were performed by the Protein Production and Structure Core Facility at EPFL. For purification, the cell pellet harvested from 300 mL of cell culture was suspended in a lysis buffer (100 mM KCl, 150 mM NaCl, 50 mM Tris, 10 mM HEPES at pH 7.6, 2 mM DTT, 1 mM EDTA, protease inhibitor cocktail, 0.2 mM PMSF, 0.1% IGEPAL, 20-30 U/mL DNase I, 5 mM MgCl_2_, and 5 mM CaCl_2_). The suspension was lysed by sonication at 30% amplitude (750W/20kHz) with 3/27 s on/off pulses for 5 cycles on ice. The lysate was clarified with three times centrifugations at 16,000-21,000 g for 15-30 minutes.

Twin-Strep affinity purification was performed on ice using 1 mL of Strep-Tactin Superflow high capacity resin (IBA Lifesciences, 2-1208-010-20ML) according to the standard gravity-flow protocol. The resin was first washed with 2 mL of buffer W (100 mM Tris, pH 7.6, 150 mM NaCl, 2 mM DTT, 1 mM EDTA) and preincubated in cold lysis buffer. After loading the lysate, the column was washed with 7 mL of W buffer and then eluted with 12 mL of E buffer (100 mM Tris, pH 7.6, 150 mM NaCl, 2 mM DTT, 1 mM EDTA, 10 mM desthiobiotin). Elution progress was monitored by SDS-PAGE gel electrophoresis (Figure S3A). The eluted sample was incubated overnight on ice with a 1.1 molar equivalent of Halo ligand-JF549 (Promega, GA1110) in buffer S (100 mM KCl, 150 mM NaCl, 10 mM HEPES at pH 7.6, 2 mM DTT). After size-exclusion chromatography on a Superdex 200 Increase 10/300 GL column (Cytiva, 28990944) (Figure S3A), the Halo-Sox TFs were assessed for labeling efficiency (92-99%) and for concentration (1-2 µM) (Figure S3A) by UV-Vis spectroscopy.

#### Electrophoretic mobility shift assays (EMSA)

The protein concentration was titrated from 0.8 nM to 50 nM while maintaining a constant concentration (2 nM) of fluorescently-labeled DNA substrate (Sdna6 in Table S9; Sox motif: 5’-CATTGTG-3’, fluorescent label: AF647)^2^. The sequence corresponds to a region (−483 to −454) of the human miR302 gene. Additionally, five-fold excess (10 nM) of poly(deoxyinosinic-deoxycytidylic) acid (poly dI-dC, ThermoFisher, 20148E) was added to reduce nonspecific interactions between Sox TFs and DNA^94^. The EMSA buffer consisted of 20 mM HEPES, 20 mM Tris, 1 mM DTT, 50 mM KCl, 0.5 mg/mL BSA, 0.02% Tween-20, 10% glycerol, 3.2% glucose, and 2 mM Trolox, at pH 7.5. After 30 minutes of incubation at room temperature, the reactions were separated by native PAGE using a 5% Tris base, boric acid, EDTA (TBE) gel, run in 0.5 x TBE buffer, at 120 V for 50 minutes on ice. Dissociation constants were estimated by quantifying the proportion of bound versus unbound DNA in each lane as the Sox TF concentration increased, based on AF647 fluorescence emission. To determine the ionic strength sensitivity of Sox2/Sox2a DNA binding via EMSA, NaCl was titrated from 12.5 mM to 400 mM into the mixture of 1 nM fluorescently-labeled DNA substrate (Sdna6 in Table S9; Sox motif: 5’-CATTGTG-3’), 10 nM Sox TFs in 20 mM HEPES, 20 mM Tris, 1 mM DTT, 6.25 mM KCl, 0.5 mg/mL BSA, 0.02% Tween-20, 10% glycerol, at pH 7.5. After 60 minutes of incubation at room temperature, the reactions were loaded separated by native PAGE (5% TBE gel, 0.5 x TBE buffer, 50 minutes at 120 V on ice). EC_50_ were estimated by quantifying the proportion of bound versus unbound DNA in each lane as the NaCl concentration increased, based on AF647 fluorescent emission.

#### Oligonucleotide labeling

Synthetic oligonucleotides containing amino modified C6 at their 5’-end were subjected to two rounds of ethanol precipitation prior to labeling. To initiate the labeling reaction, 10-20 nmoles of oligonucleotide were dissolved in 50 µl of 0.1 mM sodium tetraborate buffer (pH 8.5). A two molar-equivalent amount of the NHS ester-functionalized fluorophore (AF647 (Invitrogen, A20006) or XFD647 (AATbio, 1833), which are chemically identical) was then added to the solution. The mixture was incubated in the dark at room temperature and agitated at 300 rpm overnight. Followed by ethanol precipitation to eliminate unreacted dye, RP-HPLC purification was performed via RP-HPLC on an InertSustain C18 column (GL Sciences, 5020-07445) under a 20-minute gradient from 0 to 70% Solvent B (100 mM triethylammonium acetate, pH 7, mixed with acetonitrile), with a flow rate of 1 mL/min and detection at 260 and 640 nm. The labeled oligonucleotides were resuspended in Milli-Q water at a final concentration of 0.1-1 mM and stored at −20 °C.

#### Generation of naked and mononucleosome DNA

For DNA substrates shorter than 200 bp (Sdna3, Sdna4, Sdna5, Sdna6, Sdna8, in Table S9), two complementary oligonucleotides that are either biotinylated or AF647-conjugated (4-40 µM each) were annealed in 50-200 µl of 1× Phusion HF buffer. The mixture was heated to 95°C and then cooled to 20 °C at a rate of −1 °C/min, followed by a 1-hour incubation at 55 °C to ensure complete strand hybridization. For double-stranded DNA longer than 200 bp (Sdna9, Sdna10, Sdna11, Sdna16, in Table S9), PCR amplification was performed. Each 50 µl reaction in 1× Phusion HF buffer contained 100-600 nM of a fluorescently labeled primer, a 1.1-1.2 molar equivalent of a second primer, 200 µM dNTPs (NEB, N0447S), and 1-2 U of Phusion DNA polymerase (NEB, M0530S). The two primers were either conjugated with AF647 or biotinylated. Template DNA (1-5 ng) was typically used, except for Sdna4. For Sdna4, both primers were used at 3 µM to generate primer dimers during amplification. The thermocycling program included an initial denaturation at 95 °C for 1 minute, followed by 25-30 cycles of 95 °C for 30 seconds, 60-68 °C for 15 seconds, and 72 °C for 30 seconds, with a final 5-minute extension at 72 °C. 8-48 parallel reactions were conducted to increase DNA yield. Both the annealed and PCR-amplified products were then purified using QIAquick PCR purification kit (QIAGEN, 28104) and verified by native polyacrylamide gel electrophoresis.

**Table S9.**
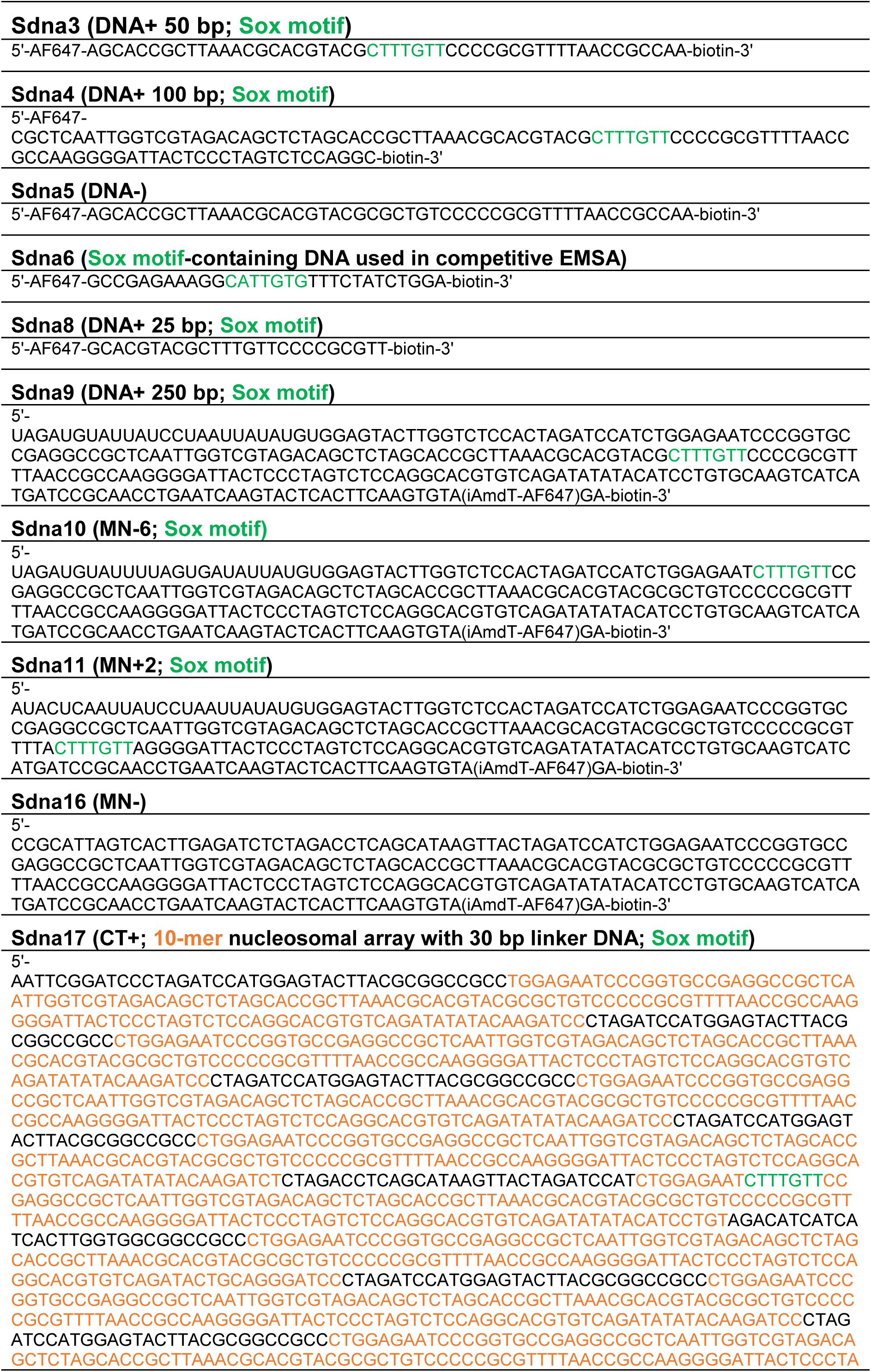
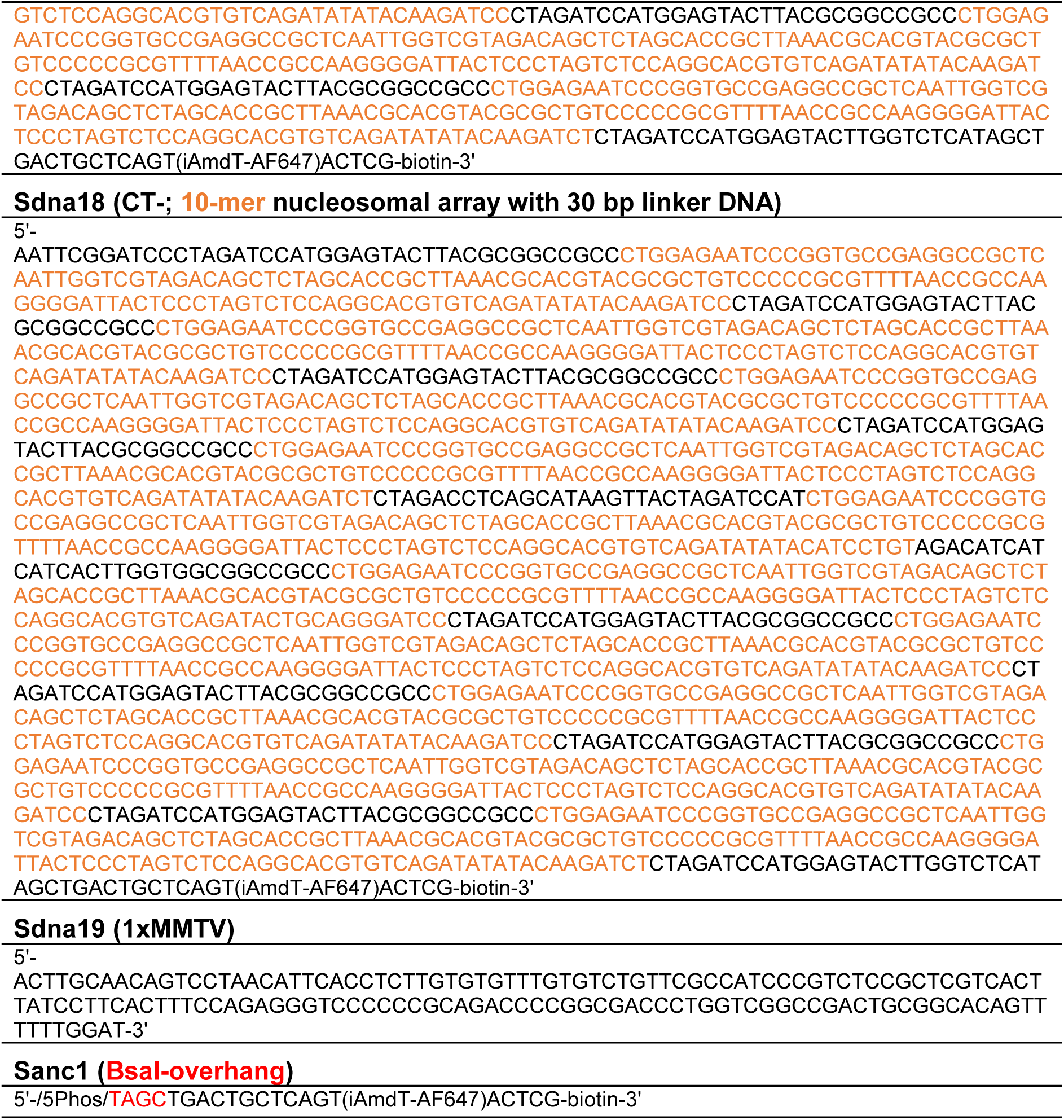
DNA sequences.

#### Preparation of 1xMMTV buffer DNA

MMTV buffer DNA (Sdna19 in Table S9) for chromatin reconstitution was derived from a plasmid carrying eight tandem repeats of the MMTV sequence (8×MMTV DNA). After growing large-scale plasmid cultures (100 mL), DNA was purified using a midiprep kit (QIAGEN, 12943) was digested overnight at 37 °C with EcoRV-HF (NEB, R3195L) and Quick CIP (NEB, M0525L), followed by incubating the digest at 80 °C for 20 minutes to inactive the enzymes. The resulting 1× MMTV fragments (151 bp) were purified from the plasmid backbone using PEG6000 precipitation. Residual PEG was removed through PCR purification kits (QIAGEN, 28104), ensuring a clean preparation of the MMTV DNA fragments.

#### Plasmid generation containing 10-mer 601WS-repeat nucleosomal array

To obtain a 10× 601WS array featuring 30 bp linkers and incorporating a Sox motif at the SHL-6 position of the fifth nucleosome (Figures 4B,S5A; Sdna17 in Table S9), we adopted a plasmid-based strategy for precision and reproducibility. The final plasmid, Spla38 was assembled through a two-step molecular cloning workflow using RecP1^95^ and Spla37. Their key sequence blocks used in this work are marked as below.

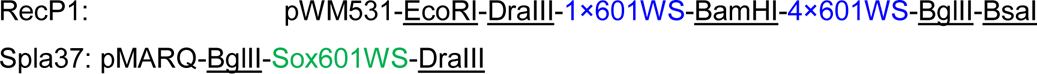

First, a plasmid (Spla42) containing 5×601WS containing a Sox motif at the fifth nucleosome was generated. To do so, **<u>BamHI</u>**-4×601WS-**<u>BglII</u>** segment from RecP1 was inserted in Spla37 (pMARQ-**<u>BglII</u>**-Sox601WS) via BamHI/BglII restriction cloning. Before digestion, Spla37 was linearized by PCR using a reverse primer (Soli179 in Table S10) containing 5’-**<u>BamHI</u>**-**<u>EcoRI</u>**-3’ restriction sites. The cloning yielded Spla42 as below.

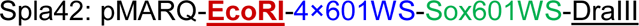

Second, a plasmid containing 10×601WS containing a Sox motif at the fifth nucleosome was cloned. The **<u>EcoRI</u>**-4×601WS-Sox601WS-**<u>DraIII</u>** segment from Spla42 was transferred back into RecP1 (pWM531-**<u>EcoRI</u>**-**<u>DraIII</u>**-5×601WS) by EcoRI/DraIII restriction cloning, which produced the final plasmid, Spla38 as below.

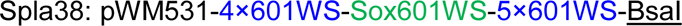

By substituting Spla37 with Spla36 (pMARQ-BglII-601WS-DraIII), we generated a plasmid (Spla35) containing 10×601WS sequence (Sdna18 in Table S9) using the same method as below.

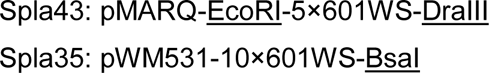

The restriction enzymes that were involved in this process: BamHI-HF, BglII, EcoRI-HF and DraIII-HF from NEB. Both Spla38 and Spla35 were validated through Sanger sequencing and restriction digest tests (EcoRI-HF, EcoRV-HF, BsaIv2-HF, and HindIII-HF; NEB). Finally, the confirmed plasmids were expanded in 200 mL cultures of DH5α E. coli and purified using the QIAquick midiprep kits (QIAGEN, 12943) followed by QIAquick PCR purification kits (QIAGEN, 28104).

#### Purification of chromatin DNA

To purify chromatin DNA for the assembly of CT+ or CT-(Sdna17 and Sdna18, respectively; sequences in Table S9), a standard digestion reaction was set up with 1000-2500 pmol of plasmid DNA (Spla38 or Spla35) in a final volume of 1.4 mL of 1× γCutSmart buffer, with addition of 1200 units of EcoRI-HF (NEB, R3101L), 2000 units of EcoRV-HF (NEB, R3195L), and 500 units of Quick CIP (NEB, M0525L). This mixture was incubated at 37 °C for 20 hours, after which the enzymes were inactivated by heating to 80 °C for 30 minutes. To make the BsaI overhang 3’-end, 1000 units of BsaI-HFv2 (NEB, R3733S) was added in a final volume of 1.46 mL of 1× γCutSmart buffer. The mixture was again incubated at 37 °C for 20 hours.

The digested DNA was subsequently purified through multiple rounds of 4-7.5% PEG6000 precipitation to eliminate the vector backbone. After adjusting the NaCl concentration to 500 mM, the sample was placed on ice for 30 minutes, and short DNA fragments were pelleted by centrifugation at 21,000 g for 30 minutes at 4 °C. Only the supernatant was retained each time, incrementally raising the PEG6000 concentration by 0.5%. Gel electrophoresis of both the supernatant and the pellet at each step monitored the purification progress. Once the desired DNA fragments were enriched to high purity, any remaining PEG was removed using QIAquick PCR purification kits (QIAGEN, 28104).

**Table S10.**
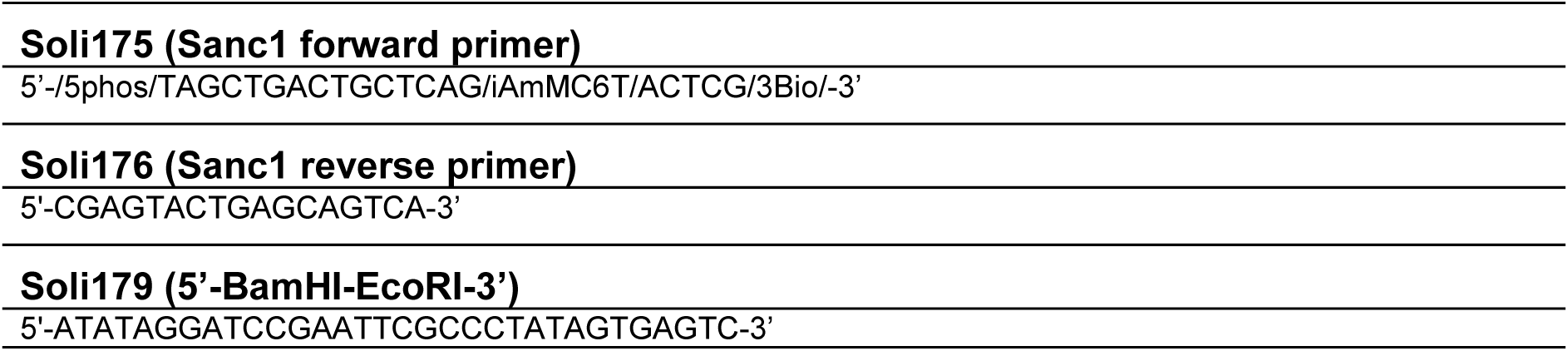
Star oligonucleotides.

A biotin-labeled oligonucleotide was annealed, to generate Sanc1 (Table S9), with its complementary strand (Soli176 and Soli175, respectively, in Table S10), which carried a 5’-phosphorylated BsaI overhang labeled via an NHS-ester AF647 protocol, in 1× T4 ligase buffer. Next, 800-1600 pmol of chromatin DNA possessing a BsaI overhang (previously purified by PEG precipitation) was mixed with a 2.5-fold molar excess of Sanc1. The mixture was then ligated overnight at 16 °C in the dark using 8,000 units of T4 DNA ligase (NEB, M0202S). Excess Sanc1 was removed by PEG precipitation, and the resulting ligated DNA (Sdna17 or Sdna18 in Table S9) was further purified with the QIAquick PCR purification kit (QIAGEN, 28104).

#### Octamer refolding

To refold histone octamers, equimolar ratios of recombinant, purified human H3.2 containing the C110A substitution and wild-type human H4 were combined in an unfolding buffer (6M Guanidine-HCl, 20 mM Tris-HCl, 5 mM DTT, pH 7.5) alongside 1.1 equivalents of wild-type human H2A and human H2B. The final protein concentration was adjusted to 0.5-1 mg/mL. Refolding proceeded via dialysis in a Slide-A-Lyzer Dialysis Cassette (ThermoFisher, 66332) against a refolding buffer (2 M NaCl, 10 mM Tris, 1 mM EDTA, 1 mM DTT, pH 7.5). Any aggregates were removed by centrifugation at 2,500 g for 20 minutes at 4 °C. The resulting octamers were purified by size-exclusion chromatography on a Superdex 200 Increase 10/300 GL column (Cytiva, 28990944) and analyzed by SDS-PAGE and RP-HPLC. Fractions containing pure octamers were pooled and concentrated to approximately 40 µM. Glycerol was then added to a final concentration of 50%, and these octamer stocks were stored at −20 °C until use.

#### Nucleosome assembly

Mononucleosomes (MN-6, MN+2, and MN-) and chromatin fibers (CT+ and CT-) containing up to 10× 601WS nucleosome positioning sequences (NPSs) with 30 bp linkers were assembled through buffer exchange to reduce the NaCl concentration from 2 M to 10 mM by dialysis (Figure S5A). In a high-salt buffer (10 mM Tris, 1 mM EDTA, 2 M NaCl), histone octamers were mixed with DNA (Sdna10, Sdna11, Sdna16, Sdna17, or Sdna18 in Table S9) at NPS-to-octamer ratios ranging from 1:1 to 1:2.5. To avoid excessive octamer binding to the fibers, 0.25 equivalents of MMTV DNA, which has lower affinity for mononucleosome formation compared to 1× 601 DNA, were added particularly to the chromatin assembly reaction. The mixture was dialyzed overnight at 4 °C, while the buffer concentration was gradually changed using a peristaltic pump from high-salt to low-salt buffer (10 mM Tris, 1 mM EDTA, 10 mM NaCl). After dialysis, any aggregates were removed by centrifugation (21,000 g, 10 minutes, 4 °C). The dialyzed sample concentrations were determined by UV using a NanoDrop spectrophotometer (ThermoFisher), and nucleosome assembly efficiency was evaluated by native PAGE on ice.

The integrity of the chromatin fibers was assessed by ScaI digestion of the fibers, resulting in individual mononucleosomes (Figure S5B). 80 ng of chromatin fibers were incubated with ScaI-HF (NEB, R3122S) at 37 °C for 3 hours in the dark. The digested products were then analyzed by native PAGE to confirm nucleosome quality and ensure full nucleosome occupancy as inferred from the absence of free DNA (Figure S5B). Once octamer saturation was verified, magnesium precipitation using 6 mM MgCl_2_ was used to further purify the chromatin fibers (Figure S5C).

#### Slide and cover glass cleaning and flow-channel assembly

Coverslips (24 × 40 mm^2^, 1.5 mm thick) and glass slides (76 × 26 mm^2^), containing holes to access flow channels after assembly, underwent multiple cleaning steps: First, they were sonicated for 20 minutes in a 10% (w/v) Alconox detergent solution, followed by successive sonication in ethanol and acetone. Between each solvent step, Milli-Q water sonication ensured removal of the previous solvent. The slides were then cleaned for 2-4 hours in a Piranha solution (3 M sulfuric acid and 60% hydrogen peroxide at a 3:1 ratio) and thoroughly rinsed in Milli-Q water. Next, they were sonicated in acetone for 10 minutes, then silanized for 10 minutes in 2% (3-aminopropyl)triethoxysilane prepared in acetone. Afterward, they were quenched in Milli-Q water and dried under nitrogen gas.

To assemble flow channels, we cut channel spacers from a 0.12 mm thickness double-sided adhesive sheet (GraceBio, 620001) using previously reported designs^24,95,96^. Each set of prepared coverslips and slides was assembled into a “sandwich” configuration, with the printed adhesive sheet sandwiched between the silanized coverslip and glass slide. The assembled devices were subsequently vacuum-sealed and stored at −20 °C until use.

#### Microfluidic channel preparation

To prepare measurement chambers, assembled coverslip-microscopy slide devices were warmed to room temperature. 10 µl pipet tips were glued into the prepared holes using epoxy glue. The individual flow-channels were then passivated by PEGylation: A PEG solution was prepared by dissolving 1 mg of biotinylated PEG-bis-Succinimidyl Valerate (SVA) (5 kDa) and 20 mg of methoxy PEG-SVA (5 kDa) in 200 µl of 100 mM sodium tetraborate buffer (pH 8.5). This solution was injected into the pre-formed channels and incubated for 2-3 hours at room temperature before use in single-molecule experiments.

#### smTIRF microscopy setup

For single-molecule colocalization imaging, we used a Nikon Ti-E inverted microscope equipped with a Nikon TIRF illuminator and a 100× oil immersion objective (NA 1.49). For illumination, two lasers were used: a Coherent OBIS 640LX (640 nm, 40 mW) to image AF647-labeled DNA and an OBIS 532LS (532 nm, 50 mW) to image JF549-labeled Halo-Sox TFs. These lasers were coupled into the microscope via an optical fiber, and an acousto-optic tunable filter was used for wavelength selection. Laser power densities at the objective were 44 W/cm^2^ for 532 nm and 11 W/cm^2^ for 640 nm. Nikon’s Perfect Focus system was used to compensate for axial fluctuations in the field of view (FOV). Emissions were detected over an 800 × 800 px² FOV by a Photometrics Prime 95B sCMOS camera, with a physical pixel size of 11 × 11 μm². Some movies with bare DNA substrates were recorded using an ANDOR iXon EMCCD camera with a 256 × 256 px² FOV and 16 × 16 μm² pixel size.

#### In vitro single-molecule colocalization measurements

To perform an experiment, all buffers were degassed before use. An oxygen-scavenging system was prepared by dissolving 1 mg catalase in 100 µl of 50 mM phosphate buffer (pH 7.0), and then by dissolving 10 mg glucose oxidase in a mixture of 40 µl catalase solution and 60 µl of T50 buffer (Gloxy). This mixture was centrifuged at 21,000 g for 3 minutes, and the supernatant was taken for use. All DNA and protein samples used for in vitro SMT were spun down at 21,000 g for 10 minutes at 4 °C to clear aggregates and were transferred to DNA low-binding (Eppendorf, 0030108051) or protein low-binding tubes (Eppendorf, 0030108116), respectively.

Prior to imaging, each PEGylated flow-channel was rinsed by ultra-pure water (ROMIL, H950L) and equilibrated with T50 buffer (10 mM Tris, 50 mM NaCl, pH 8). A peristaltic pump was used for the injection into the microfluidic channel. 30 μl of 0.2 mg/mL neutravidin in the T50 buffer was injected to facilitate DNA-substrate tethering and was washed with the T50 buffer. BSA (5 mg/mL) was incubated for 10 minutes, followed by washing with T50. Finally, the channels were equilibrated with imaging buffer (IB3) which included 20 mM HEPES, 20 mM Tris, 1 mM DTT, 3 mM EDTA, 100 mM KCl, 2 mg/mL BSA, 0.02% Tween-20, 10% glycerol, 3.2% glucose, Gloxy, and 2 mM Trolox at pH 7.5.

For immobilization, 10-50 pM of AF647-labelled, biotin-conjugated DNA constructs, mononucleosome or chromatin fibers in IB3 buffer were injected into the channel, with the degree of immobilization measured by single-molecule imaging. Excess DNA constructs, mononucleosome or chromatin fibers were washed away with 200 μl of IB3 buffer. Finally, 2 nM of JF549-labelled Halo-Sox2 or Halo-Sox2a was injected and flowed consistently at 0.5 μl/min.

After injection, Sox TF interaction dynamics with immobilized DNA, nucleosome and chromatin substrates were determined via single-molecule colocalization imaging. We recorded 10-minute movies using a 200 ms framerate, AF647-labelled DNA was detected once every 200 frames, while JF549-labelled Halo-Sox TFs were recorded every 200 ms with 100 ms illumination time and 100 ms dark time. The DNA frames were used for drift correction and localization of DNA loci, with the first DNA frame captured at 200 ms intervals.

#### ChIP-seq

Under the mESC culture conditions described above, SES4.1, SES5.1, SES7.1, and SES9.1 cells (Table S5) were incubated for 20 hours in the presence of 300, 70, 1000, and 1000 ng/mL dox, respectively, to induce expression of Halo-conjugated Sox2, Sox2a, Sox17, and Sox17b. The Halo-Sox TFs were then subsequently labeled with 50 nM Halo-SiR dye. For the ChIP experiments, two biological replicates were performed for Sox2 and Sox2a, while Sox17 and Sox17b had one replicate each.

Fixation was carried out using a PBS-based buffer, beginning with fixation at room temperature with 2 mM of disuccinimidyl glutarate (ThermoFisher, 20593) for 50 minutes and in 1% formaldehyde (ThermoFisher, 28908) for 10 minutes. The reaction was quenched by incubating the samples in 200 mM Tris-HCl (pH 8.0) for 10 minutes. Fixed cells were spun down by centrifugation at 600 g for 4 minutes at 4°C, and then were resuspended in 1% ESC-qualified fetal bovine serum (Gibco, 16141-079). The cells were sorted by FACS at 4°C, using the intensity gate resulting in the same mean intensity between SES4.1 and SES5.1 FACS samples (Figure 6A), as well as SES7.1 and SES9.1 FAC samples.

Nuclei were extracted by resuspending sorted cells in LB1 buffer (50 mM HEPES-KOH pH 7.4, 140 mM NaCl, 1 mM EDTA, 0.5 mM EGTA, 10% Glycerol, 0.5% NP-40, 0.25% Triton X-100, supplemented with Protease inhibitor cocktail (Sigma, P8340-1ML) at 1:100 dilution) and incubated for 10 minutes on ice, followed by centrifugation at 1,700 g for 5 minutes at 4 °C. This step was repeated once more. The pellets were resuspended in LB2 buffer (10 mM Tris-HCl pH 8.0, 200 mM NaCl, 1 mM EDTA, 0.5 mM EGTA, supplemented with Protease inhibitor cocktail (Sigma, P8340-1ML) at 1:100 dilution) and incubated for 10 minutes on ice, followed by the same centrifugation step. The resulting pellets were rinsed twice, without disturbing the pellets, with SDS shearing buffer (10 mM Tris-HCl pH 8.0, 1 mM EDTA, 0.15% SDS, supplemented with Protease inhibitor cocktail (Sigma, P8340-1ML) at 1:100 dilution)). The pellets were finally resuspended in SDS shearing buffer on ice and sonicated using a Covaris E220 (200 cycles, 5% duty cycle, 140 W for 20 minutes). After sonication, the samples were centrifuged at 10,000 g for 5 minutes at 4 °C to collect the supernatant.

ChIP and DNA purification were performed using ChIP-IT High Sensitivity kit (Active motif, #53040), following manufacturer instructions. Input chromatin samples were 20 μg of chromatin for Sox2 and Sox2a, and 30 μg for Sox17 and Sox17b. Drosophila spike-in chromatin (Active motif, 53083) was also added for 1 ng per 1 μg of target chromatin. Spike-in antibody (anti-H2Av; 0.5 μg per 10 ng of spike-in chromatin; Active motif, 61686) was processed together with 40 μL of Halo-Trap agarose beads (ChromoTek, ota-20) per reaction. ChIP reactions were incubated overnight at 4°C on a rotating platform at 30 rpm. Libraries were prepared with the NEBNext Ultra II DNA Library Prep Kit (NEB, E7645). The size-selected libraries using AMPure XP beads (Beckman Coulter, A63880) sequenced on an Illumina NextSeq 500 using 75-nucleotide read length paired-end sequencing.

#### ATAC-seq

SES4.1 and SES5.1 were treated with dox in the same condition as ChIP-seq, followed by labeling with 50 nM of Halo-SiR647 dyes. Each cell line was FACS-sorted at 4°C, maintaining the same intensity gate as in ChIP-seq preparation.

The ATAC-seq protocol followed established methods^97^. Approximately 50,000 cells were lysed in 50 μl of ATAC-lysis buffer (10 mM Tris-HCl, 10 mM NaCl, 3 mM MgCl_2_, 0.1% NP-40, pH 7.4), and nuclei were pelleted by centrifugation at 800 g for 5 minutes. The transposition reaction was performed by resuspending the nuclei in 50 μl of TAPS-DMF buffer (10 mM TAPS-NaOH, 5 mM MgCl_2_, 10% DMF) with addition of 0.5 μM Tn5 transposase (in-house preparation^98^), and incubating them at 37 °C for 30 minutes. DNA was purified in 10 μl of nuclease-free water, using the DNA Clean & Concentrator kit (Zymo Research, D4004) with a 5:1 ratio of binding buffer to sample.

For library preparation, the transposed DNA was PCR-amplified using NEBNext High-Fidelity 2× PCR Master Mix (NEB, M0541L) with 0.5 μM Ad1.1 universal primer, 0.5 μM Ad2.X indexing primer, and 0.6× SYBR Green I (ThermoFisher, S7585) in a total volume of 65 μl. The thermocycling program began with incubation at 72 °C for 5 minutes and 98 °C for 30 seconds, followed by 5 cycles of 98 °C for 10 seconds, 63 °C for 30 seconds, and 72 °C for 60 seconds.

10 μl of amplified DNA was analyzed by qPCR to optimize the total number of cycles based on the qPCR fluorescence signal at one-third of the saturation, to avoid the amplification saturation. The remaining DNA was further amplified accordingly, purified using column purification (Zymo, D4004), and size-selected using AMPure XP beads (Beckman Coulter, A63880). Libraries were then sequenced on an Illumina NextSeq 500 using 75 nucleotide read-length paired-end sequencing.

### QUANTIFICATION AND STATISTICAL ANALYSIS

#### In vivo SMT imaging data processing

Nuclei were detected using the Stardist plugin in Fiji^93^. Based on the nuclei ROI, single-cell movies in the JF549 channel were cropped. Using TrackIt^45^, individual Sox TF molecules within the nuclei ROI were detected by applying a detection threshold (Table S11). These spots were then linked across consecutive frames using the nearest neighbor algorithm to construct the trajectories of Sox TF molecules. The tracking loss probability (due to molecules moving laterally out of the imaging plane or out of focus) was set to 0.005. Considering the input parameters (Table S11) and the loss probability, the tracking radii were automatically computed for different dark time sets (Table S12). For the continuous movies with 100 Hz of frame rate, the tracking radius was set for all Sox TFs as 8.1312 pixels with respect to 20 µm^2^/s of the maximum diffusion coefficient of Sox2^52^.

**Table S11.** TrackIt input parameters for in vivo SMT.

| Time interval (ms) | Threshold | Min. trajectory length before gap frame | Min. trajectory length | Gap frame |
| --- | --- | --- | --- | --- |
| **TrackIt setting to measure residence times** |  |  |  |  |
| **120** | 0.7 | 2 | 2 | 3 |
| **240** | 0.7 | 2 | 2 | 2 |
| **480** | 0.7 | 2 | 2 | 1 |
| **960** | 0.7 | 2 | 2 | 1 |
| **TrackIt setting to measure diffusing dynamics** |  |  |  |  |
| **Continuous (100 Hz)** | 0.6 | 2 | 2 | 2 |

The tracking radii (*r*) were determined as described in Table S12 for in vivo SMT to measure residence times, with a pixel size of 110 nm, the maximum diffusion coefficient (*D*) was calculated as 0.055 µm^2^/s for Sox2, 0.088 µm^2^/s for Sox2a, 0.060 µm^2^/s for Sox17, 0.062 µm^2^/s for Sox17b, 0.11 µm^2^/s for Sox2D, and 0.1 µm^2^/s for Sox17D, using *r*^2^ = 4ᐧ*D*ᐧt where t = 0.12 s which is the shortest time interval between frames.

**Table S12.** Tracking radius for in vivo SMT.

| Time interval (ms) | Sox2 | Sox2a | Sox17 | Sox17b | Sox2D | Sox17D |
| --- | --- | --- | --- | --- | --- | --- |
| **120** | 1.4713 | 1.8631 | 1.5404 | 1.5635 | 2.082 | 2.0014 |
| **240** | 1.7306 | 2.3471 | 1.8746 | 1.8228 | 3.4648 | 2.3125 |
| **480** | 1.8516 | 2.4969 | 2.3759 | 1.8285 | 3.1191 | 3.7414 |
| **960** | 2.3471 | 2.8541 | 2.6121 | 1.7421 | 3.5109 | 2.9578 |

#### GRID analysis

GRID is restricted to superimposed exponential reactions with positive amplitudes and converts the inverse Laplace transformation into a minimization problem, exploring a discrete set of potential outcomes^47^. In the decay rate domain, 200 possible dissociation rates (𝑘) were logarithmically spaced from *log*(𝑘) = −3 to *log*(𝑘) = 0.85. The bulk GRID analysis yielded a photobleaching rate for each Sox TF. We applied the same photobleaching rate (0.028 s^−1^) to all Sox TF constructs, as it was determined by pooling all trajectories into the GRID analysis, regardless of the specific Sox TF.

For each Sox TF’s GRID data, 100 resamplings were performed. In each resampling, 80% of single cells at each time interval were randomly selected using a custom MATLAB code. A GRID spectrum was generated for each resampling, and any GRID-determined dissociation rate with an amplitude below 0.001 was excluded. The remaining rates from all 100 resamplings were combined into a relative frequency histogram (Figure 2C).

#### Diffusion coefficient distribution analysis

The mean squared displacement (MSD) was calculated according to the following equation:

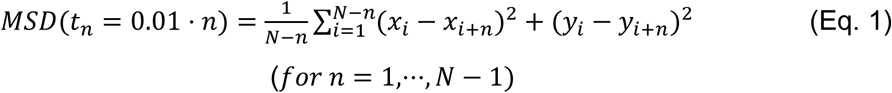

where 𝑥_𝑖_ and 𝑦_𝑖_ are the coordinates of the 𝑖-th spot within a trajectory. 𝑁 denotes the total number of spots in a given trajectory, and n represents the time lag between consecutive spots within that trajectory. The factor 0.01 reflects the 10-ms interval between movie frames.

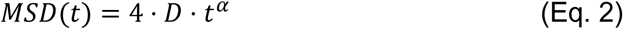

where *D* is the diffusion coefficient and 𝛼 is a coefficient to indicate the diffusion type.

Diffusion coefficient distributions were constructed fitting (Eq. 1) with (Eq. 2) using the calculation option of ‘Diffusion constants from mean square displacement fit’ in TrackIt^45^ with a power law fit on 90 % jumps per trajectory. The diffusion coefficients were then sorted per single-cell movie and binned in 200 bins logarithmically from *log*(*D*) = −2.5 to *log*(*D*) = 1.5. The bin counts were averaged across different movies for each Sox TF. The resulting mean distributions, plotted as a function of the diffusion coefficient (*D*), were then fitted with three independent Gaussian models (Eq. 3). Fitting was performed using the Levenberg-Marquardt algorithm, with each bin’s SD (from a total of 200 bins) serving as the weighting factor. The standard errors of fitting parameters were obtained from the covariance matrix of the standard errors in the fitted parameters using OriginPro 2022.

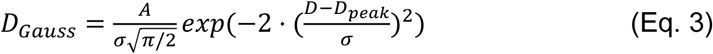

where *D_peak_* is the peak diffusion coefficient, σ is the standard deviation of the fit, and A is the area of the fit.

Figures 2D and 2E show *D_s,fit_* and *D_f,fit_*, which represent the *D_peak_* values from the respective Gaussian fits. The *F_b,fit_* in Figures 2F was calculated as below:

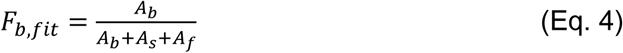

where *b*, *s*, and *f* indicate bound, slow-diffusing and fast-diffusing modes, respectively.

For statistical analysis in OriginPro 2022, the single-cell distributions of *F_b,fit_* across Sox TFs were first assessed for normality using the Shapiro-Wilk test, with normality rejected at a significance level of 0.05. Consequently, a Kruskal-Wallis ANOVA was performed, followed by Dunn’s test for multiple comparisons.

#### Spot-On analysis

The MATLAB version of Spot-On^51^ was used in this analysis. When defining the range of each diffusion coefficient in Spot-On, I referred to the intersection of the three Gaussian fits on the Sox2 diffusion coefficient distribution generated above using MSD. The ranges, in units of μm^2^/s, were set thereby as follows: 0.0005 to 0.28 for the bound state, 0.28 to 2.5 for the slow-diffusing state, and 2.5 to 25 for the fast-diffusing state. To generate the Spot-on data from single-cell analysis, the entire trajectories were used without any truncations, as per the default settings. Additionally, for z-correction, an axial resolution of 0.7 μm was applied.

#### Pseudo on-rate analysis

The pseudo on-rate (*k_pn_*) was calculated using the following formula:

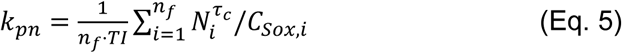

where 𝑛_𝑓_ is the number of considered frames, *TI* is a time interval between movie frames, *C_Sox,i_* is the concentration of target TFs per frame (𝑖), and 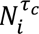 is the number of selected molecules lasting the given binding time (*τ*_𝑐_), which appear first at the frame 𝑖.

For the nonspecific *k_pn_*, we used molecules detected in 100 Hz continuous movies and *τ_c_* was set to < 1 s. To categorize bound molecules, the trajectory reconstruction process in TrackIt limited the jump distance based on a diffusion coefficient of 0.08 µm^2^/s, reflecting chromatin mobility^51,54^. Accordingly, the tracking radius was set at 0.5243 µm. The total number of trajectories—including both bound and diffusing molecules—derived from **In vivo SMT imaging data processing**, was used as a proxy for the relative concentration of Sox TFs. On the other hand, for the specific *k_pn_*, molecules detected in 120 ms-time interval movies were considered and *τ_c_* was set to > 1 s. In that case, the concentrations of Sox molecules per cell were estimated by measuring the mean intensity of Halo-JF549 within the nuclei, which was then subtracted by background mean intensity per frame. Photobleaching effects were accounted for by measuring the fluorescent intensity per frame over time.

Following the initial phase during which most molecules are detected, a series of consecutive empty frames —where no molecules are detected—could artificially lower the average molecule count, leading to an underestimation of *k_pn_*. Such empty frames often occur due to photobleaching or during long recordings with prolonged periods without signal. To correct for this bias, we defined an ‘idle time’ (5 s for nonspecific *k_pn_* and 10 s for specific *k_pn_*). When an idle period is detected, 𝑛_𝑓_ is set to the first frame of that period, and the summation for the average is terminated. This approach excludes empty frames from the calculation, ensuring a more accurate measurement of the *k_pn_*. Custom MATLAB scripts were used to perform this calculation.

For statistical analysis in OriginPro 2022, the single-cell distributions of *k_pn_* across Sox TFs were first assessed for normality using the Shapiro-Wilk test, with normality rejected at a significance level of 0.05. Consequently, a Kruskal-Wallis ANOVA was performed, followed by Dunn’s test for multiple comparisons.

#### In vitro single-molecule colocalization data processing

ND2 raw images were converted to TIFF format, followed by background subtraction using a rolling ball filter with a radius of 50 pixels. This process was done in Fiji^93^. Further data extraction was performed using custom MATLAB codes. The analysis consisted of six main steps. First, the loci of anchored DNA/MN/chromatin on the FOV were identified. Second, colocalization of Sox TFs with immobilized DNA/MN/chromatin was determined using 2D-Gaussian fitting of detected single-molecule emitters to determine the shape of the point spread function (PSF) as well as the exact position of each detection. Third, we extracted fluorescence time-traces from each DNA/MN/chromatin location. Fourth, each time-trace was segmented based on the information from the Sox TF channel (JF549 emission), i.e. sections of the trace were excluded from the analysis: Detections where the PSF exceeded 300 nm at its full width at half maximum (FWHM) were excluded. Moreover, JF549 detections that did not colocalize with DNA/MN/chromatin positions (as determined by a 300 nm threshold) were excluded from the traces. Detections that showed multiple stepwise changes in fluorescence intensity, or exceeded a threshold value were excluded from the trace, as they arose from multiple TFs or aggregates. Finally, if a trace did not show at least two TF binding detections, the trace was eliminated from the set. If more than 80% of traces from a movie were excluded, the whole movie was discarded. Fifth, traces were fitted using a stepfit algorithm^99^. Finally, bound states were identified using an intensity threshold from the fitted trace and bound and unbound times were determined and saved.

#### Statistical analysis for residence times, on-rates and specific on-rate

The bound, unbound, and search times from each movie (Figure 3E) were used to generate the corresponding 1-CDF curves (Figures 3G, S3D, S3I). First, a cumulative histogram was constructed with a bin size of 0.2 s—matching the movie frame interval. Then, the histogram was normalized to range from 0 to 1. To estimate the residence times of Sox TFs, we applied multi-exponential decay fits to the 1-CDF curves of bound times. For naked DNA substrates, a biexponential model (n=2 in Eq. 6) was used, while for nucleosomes, a triexponential model (n=3 in Eq. 6) was employed.

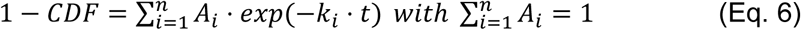

In these models, the inverse of each decay rate, *k_i_*, corresponds to a residence time, *τ_i_*. In contrast, on-rate (*k_on_*) and specific on-rate (*k_s,on_*) were determined by fitting the 1-CDF curves of unbound times with a monoexponential model (n=1 in Eq. 6); here, the *k_on_* and *k_s,on_* were calculated by dividing the decay rate *k_1_* by the Sox TF concentration (2 nM). All fitting procedures were carried out using the Levenberg–Marquardt algorithm in OriginPro 2022.

#### DNA coiling model

DNA, as a polymer, will coil when its length exceeds its persistence length (𝑙_𝑝_), which is approximately 45 nm^57–59^—roughly equivalent to 132 bp. For naked DNA of length 2𝐿, the gyration radius (𝑅_𝑔_) is expressed as follows:

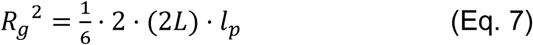

The on-rate (*k_on_*) may depend on the overall size of the DNA encountered by a TF. Because DNA coils, *k_on_* is proportional to the DNA’s gyration radius (𝑅𝑅_𝑔𝑔_) multiplied by the nanoscopic association rate (*k_a_*), which is defined as the rate at which a TF binds to a unit-length DNA segment (2𝐿_0_).

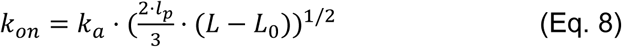

Nonlinear fitting of the *k_on_* curve (Figure S3F) using Eq. 8 was performed with the Levenberg– Marquardt algorithm, employing the standard deviation of each data point as the weighting factor. The standard errors of the fitting parameters *k_a_* and 𝐿_0_ were calculated with OriginPro 2022. In this fitting, the persistence length (𝑙_𝑝_) was fixed at 132 bp. 𝐿_0_ was assumed to be the same between Sox2 and Sox2a, which was obtained as 5.6 bp. Statistical significance in *k_a_* (Figure S3H) was determined using a two-tailed t-test, with the fitted parameter values as the mean, their standard errors as the standard deviation, and four as the sample size.

#### 1D target search model

A 1D search model has been previously introduced by Berg et al.^11^ and Hammar et al.^15^. It describes the behavior of a TF on bare DNA containing a single binding site at its center. We can use this model to describe the Sox2 target search for a motif located at the center of a stretch of naked DNA (Figure 3I). The model involves four parameters (Table S5): the nanoscopic association rate (*k_a_*), nonspecific residence time (*τ_d_*), the 1D diffusion coefficient (*D_1D_*), and the target recognition rate (*k_r_*). A TF binds to any sites on DNA at *k_a_*, slides with *D_1D_* and then recognizes the target with *k_r_* upon encounter. During this process, the TF resides on DNA nonspecifically during *τ_d_*.

In this model, the specific on-rate (*k_s,on_*) was defined as the association rate to the target site on bare DNA, via 1D diffusion as well as 3D diffusion. Consequently, *k_s,on_* depends on *L*, the length of flanking DNA as below:

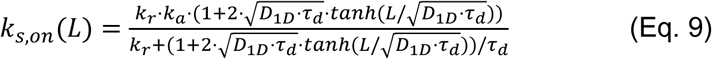

Although *k_r_* cannot be measured directly at present, it can be indirectly estimated (Figure 3M) by determining the specific on-rate (*k_s,on_*) in a defined system under various conditions (Figure 3J).

To fit the measured *k_s,on_* curve (Figure 3J) with the 1D target search model, we considered the sliding length (𝑠𝑠_𝐿𝐿_), which was previously introduced as below:

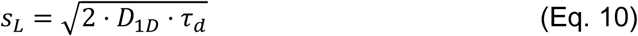

Furthermore, we defined a target recognition constant (𝐾𝐾_𝑅𝑅_) as below, since the target recognition process would occur before it dissociates from the site:

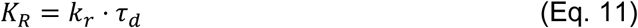

Using Eq. 10 and Eq. 11, *k_s,on_* in Eq. 9 becomes as a function of *L* with three parameters of *k_a_*, *K_R_*, and *s_L_*, as below:

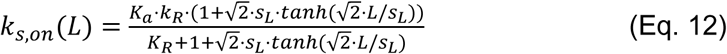

Accordingly, the recognition probability *p_bind_* is expressed as below:

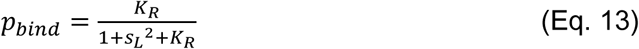

The nonlinear fitting on the *k_s,on_* curve (Figure 3J) with Eq. 12 was conducted based on the Levenberg-Marquardt algorithm using the SD of each data point as the weighting factor. The standard errors of fitting parameters or derived parameters of *k_r_* and *D_1D_* (from Eq. 10 and Eq. 11 considering *τ_d_* = 10 ms^61^) were also calculated in OriginPro 2022. In the fitting, *k_a_* was fixed to measured values of *k_a,Sox2_* = 2.13 M^−1^s^−1^bp^−1^ and *k_a,Sox2a_* = 1.91 10^5^ M^−1^s^−1^bp^−1^. The statistical significance was determined using a two-tailed t-test, with the fitted parameter values as the mean, their standard errors as the standard deviation, and four as the sample size.

#### ChIP-seq analysis

ChIP-sequencing libraries were mapped (BAM format) to mm10 version of Mus musculus genome or version BDGP6 of Drosophila Melanogaster genome, using STAR 96 with parameters --alignMatesGapMax 2000 --alignIntronMax 1 --alignEndsType EndtoEnd -- outFilterMultimapNmax 1. Duplicated reads and reads not mapping to chromosome 1-19 and X or Y, were removed using SAMTools^100^. For the generated Drosophila alignments, the number of reads was determined, and normalization factors were calculated to bring all the Drosophila bam files to the same number of reads. These normalization factors were then applied to downsample alignment files mapped on the mouse genome using SAMTools^100^. Besides, between two replicates (i.e. for Sox2 and Sox2a), the ratio of their sequencing depths were used as a normalization factor, which were applied to downsample alignment files mapped on the mouse genome using SAMTools^100^. Peak calling was then performed on these downsampled bam files using MACS2 v 3.0.0a7 with settings -f BAMPE -g mm. The regions contained in the mm10 blacklist (ENCODE consortium) were excluded. For the score comparison between different samples, the bam files of replicates were merged using SAMTools^100^. Scores were determined from downsampled or merged bam files using the bamCoverage function from DeepTools v3.3.0^101^ with setting -- normalizeUsing RPKM.

Between replicates, ChIP peaks were consolidated using a specific combining protocol (https://ro-che.info/articles/2018-07-11-chip-seq-consensus), employing lower=1 and rangesOnly=T in the slice() function from the GenomicRanges library. Next, these merged peak files were used to generate Sox-enriched regions following the same procedure. Starting from the replicate-merge bed files, CORs were defined via the same protocol, but with lower=2 and rangesOnly=T in slice(). To quantify enrichment across CORs, two approaches were used: (i) multiBigwigSummary in DeepTools^101^ or (ii) bigWigToBedGraph from the UCSC genome toolkit. Log2-fold changes in Sox2 scores relative to Sox2a scores were computed for each COR, and regions were classified as Sox2-enriched or Sox2a-enriched based on the sign of the log2-fold values. For exclusive region analysis, each replicate-merged bed file (Sox2 or Sox2a) was processed with the subtract function in BEDTools^102^ to remove CORs. The resulting intervals were then merged again using the same ChIP peak consolidation protocol.

Next, these replicate-merged bed files were input into HOMER2’s findMotifsGenome.pl^103^ to discover motifs. The search was restricted to 200 bp regions around the peaks, on bigWig files that were generated from merged BAM files spanning the replicates.

#### ATAC-seq analysis

ATAC-Seq libraries were mapped and filtered as described for ChIP-Seq. Accessible peaks were then called, and scores were computed as described above for ChIP-Seq.

#### Statistical analysis

Most results are presented as means ± standard deviation (SD) unless stated otherwise. Two-group comparisons were conducted using two-tailed, heteroscedastic t-tests. For multiple comparisons, distributions were first evaluated for normality using the Shapiro-Wilk test; since normality was rejected in all cases, a nonparametric, one-way Kruskal-Wallis ANOVA was performed, followed by Dunn’s test for pairwise comparisons. A p-value < 0.05 was considered statistically significant.

## DATA AND CODE AVAILABILITY

Microscopy data, evaluation and analysis scripts, and detailed plasmid maps of expression vectors are available upon request.

All single-molecule data is available from Zenodo (currently not public): https://doi.org/10.5281/zenodo.14948339 (Figures 1, S1); https://doi.org/10.5281/zenodo.14948341 (Figures 2, S2); https://doi.org/10.5281/zenodo.14948309 (Figures 3, S3); https://doi.org/10.5281/zenodo.14948299 (Figures 4-5, S4-S5).

All sequencing data were deposited at NCBI Gene Expression Omnibus (GEO), accession number GSE290873 and GSE290874 (accessible with private tokens): https://www.ncbi.nlm.nih.gov/geo/query/acc.cgi?acc=GSE290873 https://www.ncbi.nlm.nih.gov/geo/query/acc.cgi?acc=GSE290874

## Notes

### Competing Interest Statement

The authors have declared no competing interest.

