## Supplementary Figures and Tables 1-4 for "Electrostatic properties of disordered regions control transcription factor search and pioneer activity"

#### SUPPLEMENTARY INFORMATION

##### Supplementary Figures

**Figure S1**

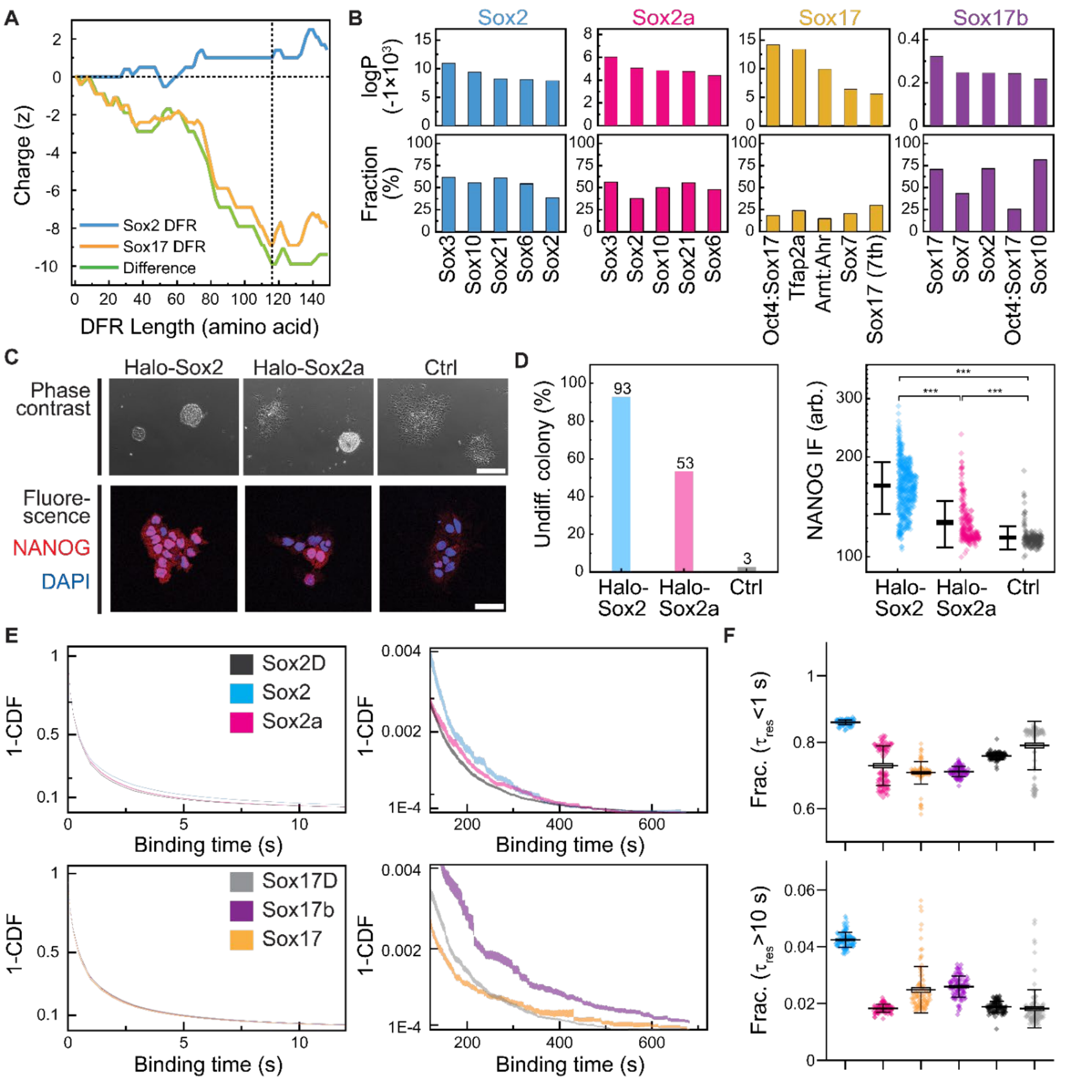


**Figure S1. Characterization of biophysical and functional properties of Sox TFs**

**(A)** Net charge curves of the C-terminal DFRs of Sox2 and Sox17, and the difference between them as a function of the accumulated length in amino acids. The horizontal dashed line indicates where the net charge is zero, and the vertical dashed line marks the point where the charge difference is maximized while minimizing the DFR length.

**(B)** HOMER2 known motif analysis on ChIP-seq data for mESCs overexpressing Sox TFs. The Top 5 ranked motifs are listed.

**(C)** Representative images of phase-contrast and NANOG immunofluorescence (IF) staining for 2TS22C cells expressing Halo-Sox2 (left), Halo-Sox2a (middle) or control 2TS22C cells (no transgene of Halo-Sox TFs), with dox addition.

**(D)** Left: Quantification of sharply-edged, dome-shaped colonies representing an undifferentiated state, based on phase contrast images (N = 56, 92, 37 cells). Right: Single-cell analysis of NANOG IF intensity. Thick black line: Mean. Error bar: SD (N = 487, 232, 184 cells).

**(E)** Cumulative distribution of binding times for Sox TFs in two distinct temporal regimes (short and long), corresponding to a zoomed-in view of Figure 1E. Curve shading: SD.

**(F)** Bound fractions of Sox TFs that last shorter than 1 s (top) and longer than 10 s (bottom). Each point represents each resampled data set (N=100). Thick black line: Mean. Box height: SE. Error bar: SD.

#### Figure S2

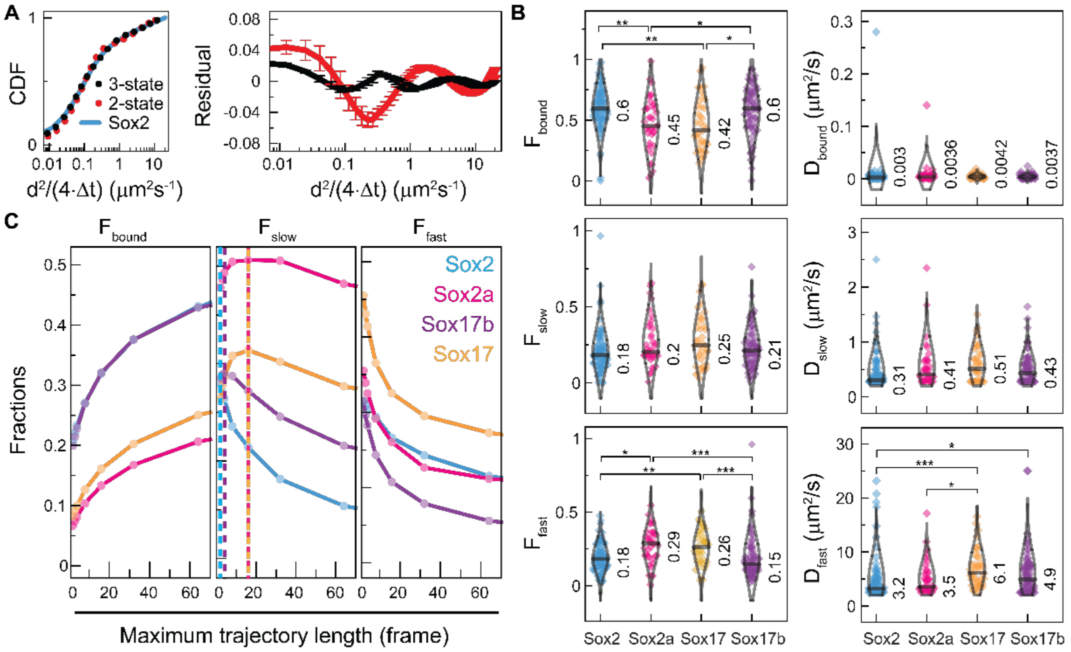


##### Figure S2. Sox TFs with $DFR_{Sox2}$ rapidly transit from slow diffusing to bound state.

**(A)** Left: cumulative jump distance (d) distributions of Sox2 (blue), fitted with a biexponential (two-state model; red) and a triexponential (three-state model; black) curve, considering $\Delta t$ = 10 ms. Right: Residual curves of the two multiexponential fits. Each curve was averaged across Sox2, Sox2a, Sox17, and Sox17b. Cap: SD.

**(B)** Spot-On results from single-cell analysis considering the three-state diffusion model (N = 86, 38, 38, 72 cells). Black line: median. Number: median value. Statistical significance was evaluated using Dunn-test following a nonparametric Kruskal-Wallis ANOVA (*: p≤0.05; **: p≤0.01; ***: p≤0.001) and non-significant comparisons are not marked.

**(C)** Spot-on fractions of bound (left), slow diffusion (center), and fast diffusion (right) modes as a function of maximum trajectory length (upper truncation limit used in the analysis). The vertical lines indicate the peak position of each curve (color-coded).

**Figure S3**

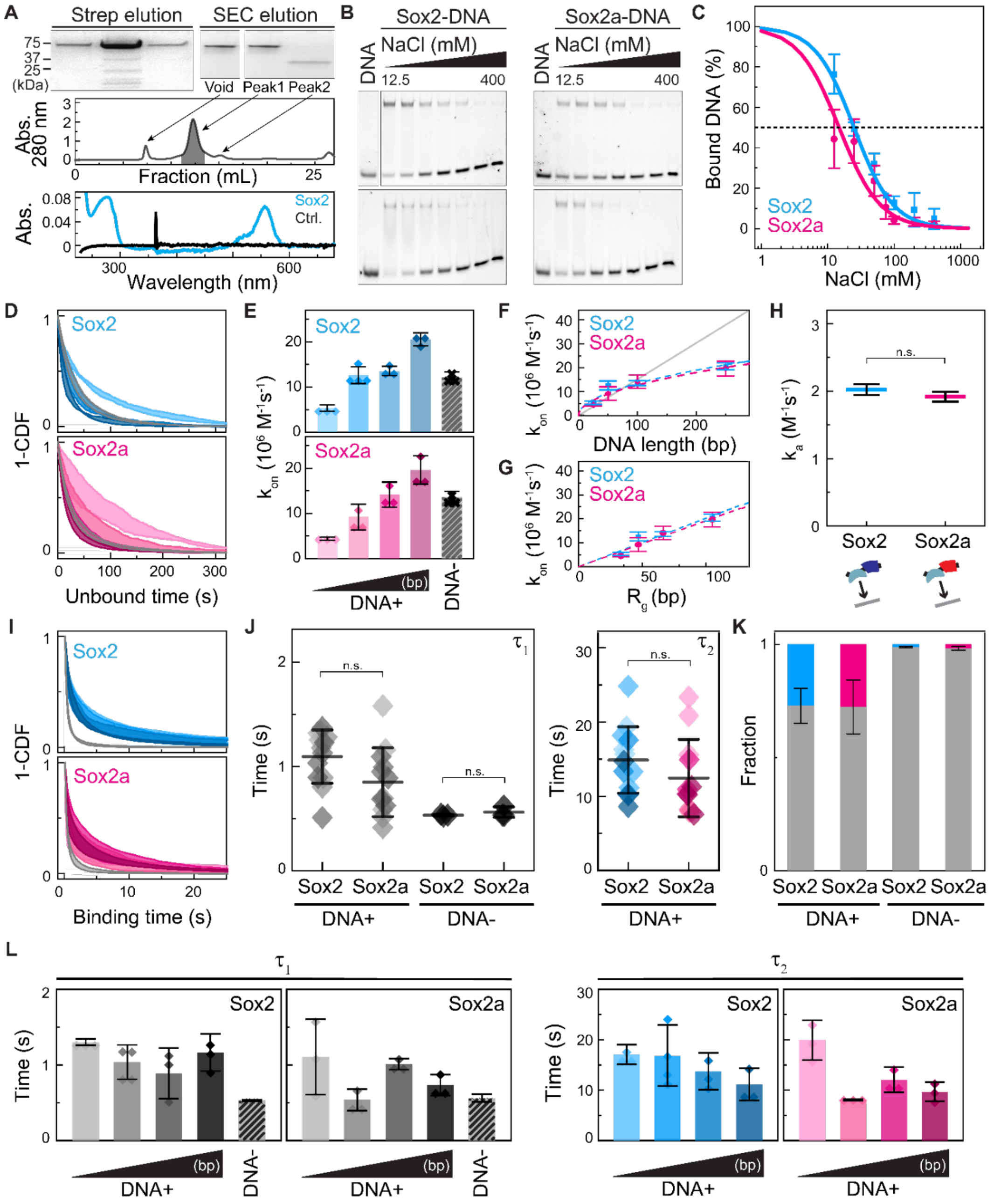


#### Figure S3. DFR charge does not affect 3D diffusion-mediated association or residence times on naked DNA

**(A)** Top: SDS-PAGE gels of Sox2 purified using Strep-tag purification (Strep elution) followed by size exclusion chromatography (SEC elution). Middle: Absorbance curve at 280 nm of the SEC elution. The filled area indicates the final collected product. Bottom: UV-Vis absorbance spectrum of purified JF549-labeled Halo-Sox2 at a two-fold dilution. The control (Ctrl.) is the S buffer **(see Methods)** used to dissolve Halo-Sox2, without proteins.

**(B)** NaCl-titrating EMSA visualized in the AF647 emission channel to determine the ionic strength of Sox-DNA complex.

**(C)** Quantification of the fraction of bound DNA from the NaCl-titrating EMSA (N=3) shown in (B) and Figure 1C. Data were fitted using the Hill equation.

**(D)** Cumulative distributions of unbound times for Sox2 (top, blue) and of Sox2a (bottom, pink) on various naked DNA substrates. Curve shading: SD.

**(E)** On-rate of Sox TFs obtained by fitting the 1-CDF curves in (D) with a monoexponential decay. Symbol: replicate (N=3-4). Bar: mean. Cap: SD.

**(F)** On-rate fit curve as a function of DNA length (dashed line), regarding the DNA polymer model. Symbol: replicate (N=3-4). Cap: SD.

**(G)** On-rate fit curve as a function of DNA gyration radius (dashed line), regarding the DNA polymer model. Symbol: replicate (N=3-4). Cap: SD.

**(H)** Nanoscopic association rate of Sox TFs, obtained from the fitting in (F) and (G). Line: mean. Cap: fitting error.

**(I)** Cumulative distributions of binding times for Sox2 (top, blue) and of Sox2a (bottom, pink) on various naked DNA substrates. Curve shading: SD.

**(J)** Short-lived ($\tau_1$) and long-lived ($\tau_2$) residence times of Sox TFs on various naked DNA, depending on the motif presence (DNA+ or DNA-). Symbol: replicate per color-coded DNA length. Bar: mean. Cap: SD.

**(K)** Corresponding bound fractions of Sox TFs for the residence times in (J). The fraction of short-lived ($\tau_1$, gray), and long-lived ($\tau_2$, colored) populations are shown in bar graphs. Bar: mean. Cap: SD.

**(L)** Residence times of Sox TFs on naked DNA of varying lengths. Symbol: replicate (N=3-4). Bar: mean. Cap: SD.

Replicates: N=4 for Sox2 with 50 bp DNA+. N=3 for all the other cases.

Statistical significance was assessed using two-tailed t-tests (*: $p \leq 0.05$; **: $p \leq 0.01$; ***: $p \leq 0.001$).

**Figure S4**

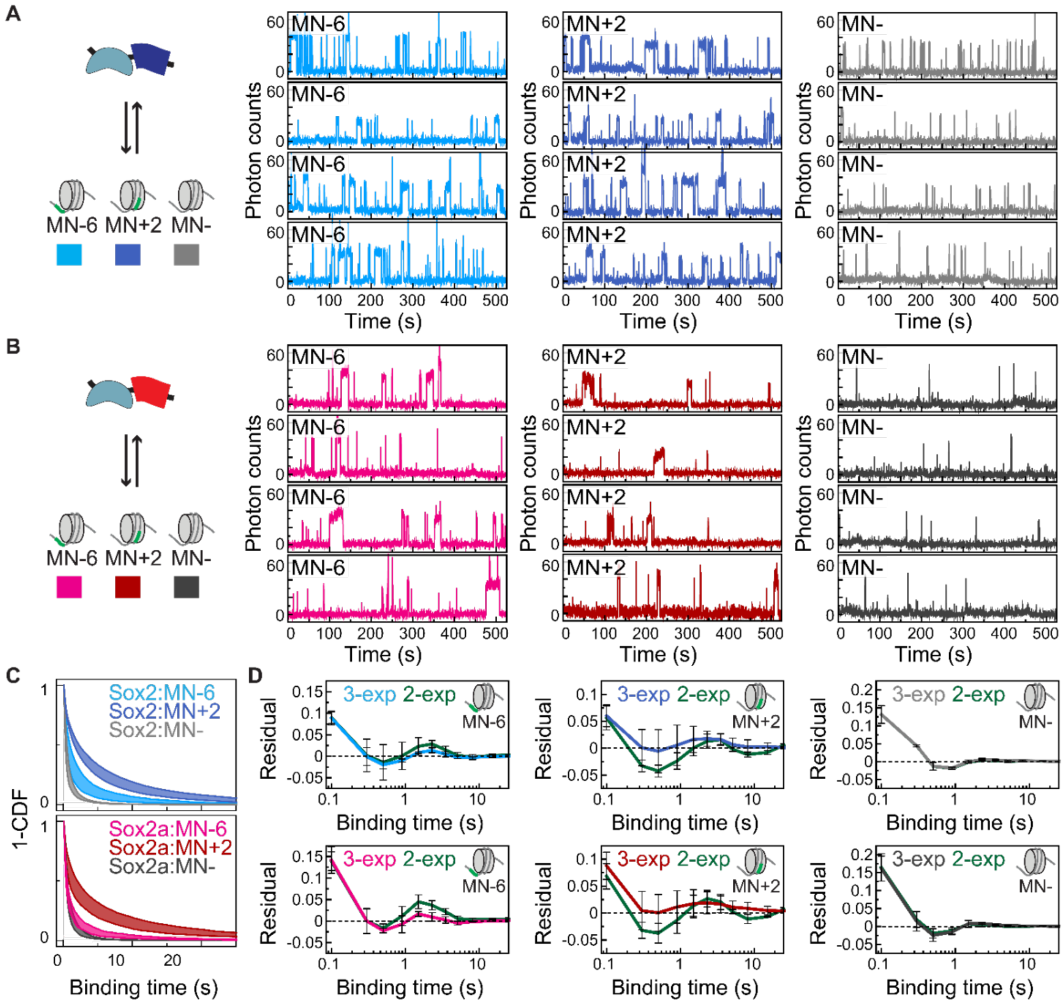


**Figure S4. Sox TFs exhibit three distinct bound states on nucleosomes.**

**(A-B)** Representative fluorescence time-traces of Sox2 **(A)** and Sox2a **(B)** interactions with different mononucleosome constructs.

**(C)** Cumulative distributions of binding times for Sox2 (top, blue) and of Sox2a (bottom, pink) on various mononucleosome constructs. Curve shading: SD.

**(D)** Residual curves of biexponential (2-exp) and triexponential (3-exp) decay fits on (C). Line: mean. Cap: SD.

Replicates: N=4 for Sox2 with MN+2 and CT+, and Sox2a with MN+2. N=3 for all the other cases.

#### Figure S5

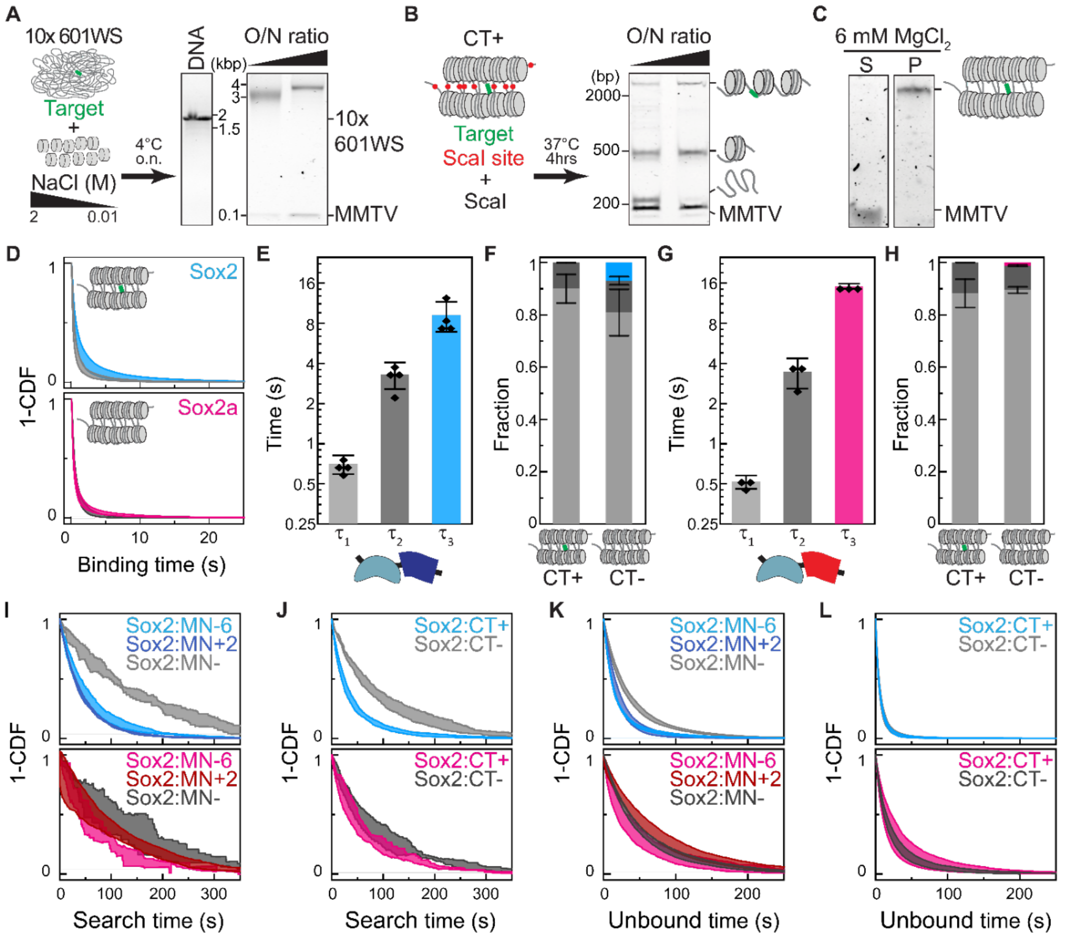


**Figure S5. DFR$_{Sox2}$ confers superior specific binding on chromatin.**

**(A)** In vitro chromatin reconstitution using NaCl-diluting dialysis. The molar ratio of octamer (O) to nucleosome positioning sequence (N) changed to optimize the octamer input concentration. Dialyzed products were loaded in 0.6 % agarose gel which was imaged using SYBR Safe.

**(B)** Validation of octamer saturation in chromatin fibers using ScaI digestion. The digested samples were loaded in 5% TBE PAGE gel which was imaged using SYBR Safe.

**(C)** $Mg^{2+}$ precipitation to purify chromatin fibers from MMTV buffer DNA that was used to compensate for overloaded octamers in chromatin assembly. Supernatant (S) and pellet (P) were loaded in 0.6 % agarose gel which was imaged using SYBR Safe.

**(D)** Cumulative distributions of binding times for Sox2 (top, blue) and of Sox2a (bottom, pink) on motif-containing (CT+) or motif-absent (CT-) chromatin fibers. Curve shading: SD.

**(E-H)** Residence times and corresponding bound fractions of Sox2 **(E-F)** and Sox2a **(G-H)** on chromatin fibers. Symbol: replicate (N=3-4). Bar: mean. Cap: SD. The bound fractions are color-matched to short-lived ($\tau_1$, gray), intermediate-lived ($\tau_2$, dark gray) and long-lived ($\tau_3$, blue) residence times.

**(I-J)** Cumulative distributions of search times for Sox2 (top, blue) and of Sox2a (bottom, pink) on mononucleosomes **(I)** and chromatin fibers **(J)**. Curve shading: SD.

**(K-L)** Cumulative distributions of unbound times for Sox2 (top, blue) and of Sox2a (bottom, pink) on mononucleosomes **(K)** and chromatin fibers **(L)**. Curve shading: SD.

Replicates: N=4 for Sox2 with MN+2 and CT+, and Sox2a with MN+2. N=3 for all the other cases.

### Figure S6

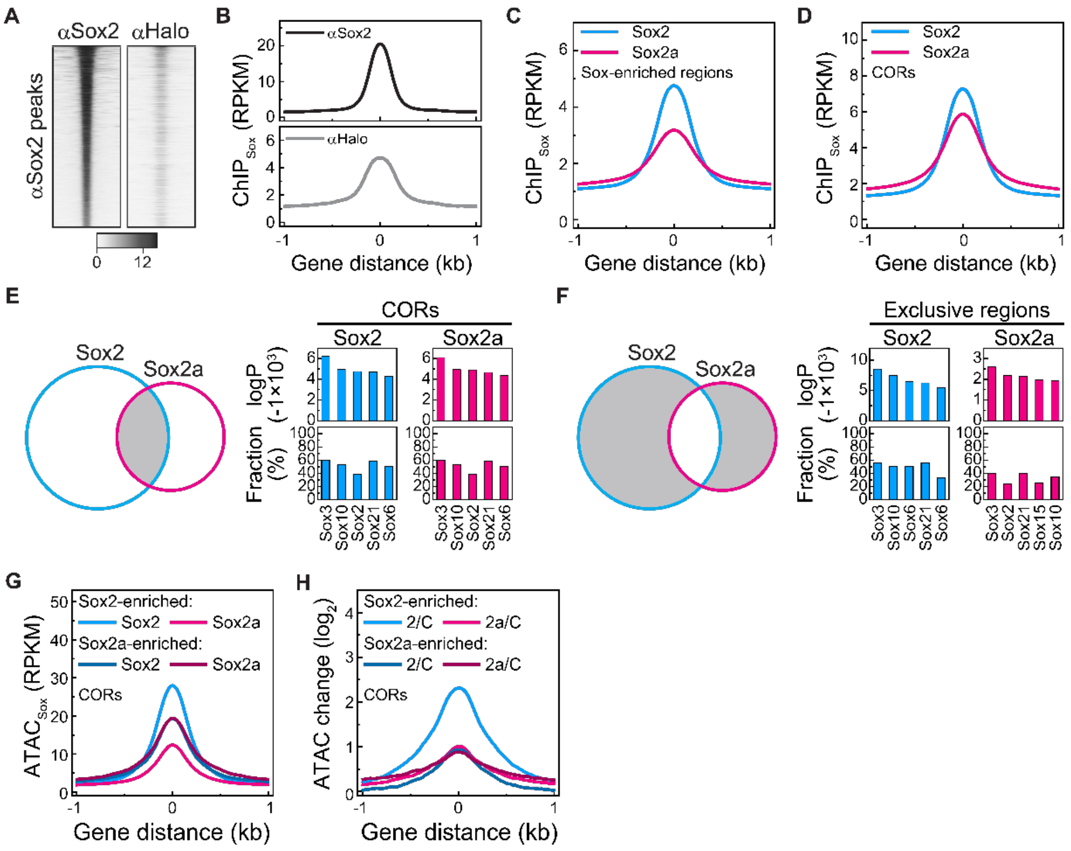


**Figure S6. DFR$_{Sox2}$ enhances pioneer activity.**

**(A)** Heatmaps of anti-Sox2 (αSox2) and anti-Halo (αHalo) ChIP-seq data (RPKM) across αSox2 ChIP peaks.

**(B)** Mean score profile of αSox2 (top) and αHalo (bottom) ChIP-seq data across anti-Sox2 ChIP peaks, corresponding to the heatmaps in (A).

**(C-D)** Mean score profile of ChIP-seq data for mESCs overexpressing Sox TFs across Sox-enriched regions **(C)** and across CORs **(D)**, defined in Figure 6C.

**(E-F)** HOMER2 known motif analysis on ChIP-seq data for mESCs overexpressing Sox TFs across CORs **(E)** and across exclusive regions **(F)**. The Top 5 ranked motifs are listed.

**(G)** Mean score profile of Sox2 and Sox2a ATAC data across Sox2-enriched and Sox2a-enriched CORs.

**(H)** Mean score profile of the log2-fold change in ATAC scores of Sox TFs over Ctrl, across Sox2-enriched and Sox2a-enriched CORs.

#### Supplementary Tables

##### Table S1

| GRID parameters | Sox2 | | Sox2a | | Sox17 | | Sox17b | | Sox2D | | Sox17D | |
|---|---|---|---|---|---|---|---|---|---|---|---|---|
| | Mean | SD | Mean | SD | Mean | SD | Mean | SD | Mean | SD | Mean | SD |
| $F_{t<1s}$ | 0.860 | 0.008 | 0.729 | 0.059 | 0.708 | 0.034 | 0.712 | 0.015 | 0.759 | 0.011 | 0.790 | 0.073 |
| $F_{t>10s}$ | 0.042 | 0.003 | 0.018 | 0.001 | 0.025 | 0.008 | 0.026 | 0.004 | 0.019 | 0.002 | 0.018 | 0.007 |
| **Diffusing parameters** | **Sox2** | | **Sox2a** | | **Sox17** | | **Sox17b** | | **Sox2D** | | **Sox17D** | |
| | Mean | SD | Mean | SD | Mean | SD | Mean | SD | Mean | SD | Mean | SD |
| $D_{b,peak}$ ($\mu m^2 s^{-1}$) | 0.138 | 1.067 | 0.173 | 1.111 | 0.181 | 1.152 | 0.131 | 1.066 | - | - | - | - |
| $D_{s,peak}$ ($\mu m^2 s^{-1}$) | 2.024 | 1.053 | 2.163 | 1.047 | 2.583 | 1.063 | 2.285 | 1.047 | - | - | - | - |
| $D_{f,peak}$ ($\mu m^2 s^{-1}$) | 7.469 | 1.057 | 7.733 | 1.041 | 8.385 | 1.033 | 8.399 | 1.033 | - | - | - | - |
| $F_{b,fit}$ | 0.629 | 0.078 | 0.514 | 0.104 | 0.458 | 0.123 | 0.652 | 0.066 | - | - | - | - |
| $F_{s,fit}$ | 0.300 | 0.127 | 0.368 | 0.143 | 0.366 | 0.168 | 0.262 | 0.107 | - | - | - | - |
| $F_{f,fit}$ | 0.071 | 0.257 | 0.119 | 0.194 | 0.176 | 0.196 | 0.086 | 0.145 | - | - | - | - |
| $F_{b,filled}$ | 0.259 | 0.127 | 0.178 | 0.120 | 0.155 | 0.113 | 0.271 | 0.150 | - | - | - | - |
| **$k_{pn}$ parameters** | **Sox2** | | **Sox2a** | | **Sox17** | | **Sox17b** | | **Sox2D** | | **Sox17D** | |
| | Mean | SD | Mean | SD | Mean | SD | Mean | SD | Mean | SD | Mean | SD |
| $k_{ns,pn}$ | 1.000 | 0.530 | 0.655 | 0.217 | 0.681 | 0.394 | 0.898 | 0.451 | - | - | - | - |
| $k_{s,pn}$ | 1.000 | 0.615 | 0.397 | 0.243 | 0.587 | 0.458 | 0.280 | 0.187 | - | - | - | - |
| **Spot-on parameters** | **Sox2** | | | **Sox2a** | | | **Sox17** | | | **Sox17b** | | |
| | Mean | SD | Med. | Mean | SD | Med. | Mean | SD | Med. | Mean | SD | Med. |
| $D_{bound}$ ($\mu m^2 s^{-1}$) | 0.007 | 0.008 | 0.005 | 0.004 | 0.030 | 0.022 | 0.004 | 0.004 | 0.003 | 0.004 | 0.004 | 0.004 |
| $D_{slow}$ ($\mu m^2 s^{-1}$) | 0.452 | 0.551 | 0.595 | 0.503 | 0.331 | 0.414 | 0.329 | 0.282 | 0.306 | 0.409 | 0.511 | 0.428 |
| $D_{fast}$ ($\mu m^2 s^{-1}$) | 4.806 | 5.051 | 6.940 | 6.038 | 4.118 | 3.153 | 3.572 | 4.260 | 3.226 | 3.549 | 6.132 | 4.930 |
| $F_{bound}$ | 0.606 | 0.174 | 0.598 | 0.464 | 0.221 | 0.451 | 0.464 | 0.213 | 0.418 | 0.580 | 0.200 | 0.596 |
| $F_{slow}$ | 0.205 | 0.139 | 0.183 | 0.261 | 0.160 | 0.204 | 0.276 | 0.170 | 0.246 | 0.231 | 0.137 | 0.212 |
| $F_{fast}$ | 0.189 | 0.096 | 0.182 | 0.274 | 0.123 | 0.288 | 0.260 | 0.120 | 0.262 | 0.188 | 0.148 | 0.148 |

**Table S1. In vitro SMT analysis of Sox TFs diffusing and binding dynamics**

Standard deviation (SD) and Median (Med.)

#### Table S2

| Data label | TF | DNA | $t_1$ ($s^{-1}$) | $t_2$ ($s^{-1}$) | $A_1$ | $A_2$ | $k_{on}$ ($M^{-1}s^{-1}$) | $k_{s,on}$ ($M^{-1}s^{-1}$) |
|---|---|---|---|---|---|---|---|---|
| **DNA01** | Sox2 | 25 bp | 1.318 | 16.303 | 0.621 | 0.379 | 5865000 | 5146698 |
| **DNA02** | Sox2 | 25 bp | 1.332 | 19.301 | 0.681 | 0.319 | 4585411 | 4245000 |
| **DNA03** | Sox2 | 25 bp | 1.253 | 15.695 | 0.759 | 0.241 | 5687951 | 5119549 |
| | | ***average*** | ***1.301*** | ***17.099*** | ***0.687*** | ***0.313*** | ***5379454*** | ***4837082*** |
| **DNA04** | Sox2 | 50 bp | 1.281 | 13.305 | 0.645 | 0.355 | 11761200 | 8478339 |
| **DNA05** | Sox2 | 50 bp | 0.803 | 18.199 | 0.859 | 0.141 | 15411800 | 9903372 |
| **DNA06** | Sox2 | 50 bp | 0.892 | 11.150 | 0.611 | 0.389 | 11949500 | 7992730 |
| **DNA07** | Sox2 | 50 bp | 1.176 | 24.815 | 0.713 | 0.288 | 11396300 | 7635883 |
| | | ***average*** | ***1.038*** | ***16.867*** | ***0.707*** | ***0.293*** | ***12629700*** | ***8502581*** |
| **DNA08** | Sox2 | 100 bp | 1.033 | 13.356 | 0.752 | 0.248 | 14762600 | 12964300 |
| **DNA09** | Sox2 | 100 bp | 1.129 | 17.575 | 0.679 | 0.321 | 13041700 | 12407500 |
| **DNA10** | Sox2 | 100 bp | 0.509 | 10.296 | 0.765 | 0.235 | 12894800 | 13645000 |
| | | ***average*** | ***0.890*** | ***13.742*** | ***0.732*** | ***0.268*** | ***13566367*** | ***13005600*** |
| **DNA11** | Sox2 | 250 bp | 0.895 | 14.717 | 0.825 | 0.175 | 18905600 | 17206500 |
| **DNA12** | Sox2 | 250 bp | 1.372 | 10.170 | 0.789 | 0.211 | 21254500 | 15119700 |
| **DNA13** | Sox2 | 250 bp | 1.228 | 8.612 | 0.771 | 0.229 | 21540000 | 14591800 |
| | | ***average*** | ***1.165*** | ***11.166*** | ***0.795*** | ***0.205*** | ***20566700*** | ***15639333*** |
| **DNA14** | Sox2 | DNA- | 0.523 | 7.201 | 0.987 | 0.013 | 11336700 | 6981426 |
| **DNA15** | Sox2 | DNA- | 0.539 | 11.933 | 0.984 | 0.016 | 11689200 | 6896051 |
| **DNA16** | Sox2 | DNA- | 0.531 | 10.646 | 0.991 | 0.009 | 13512700 | 5913263 |
| | | ***average*** | ***0.531*** | ***9.927*** | ***0.987*** | ***0.013*** | ***12179533*** | ***6596913*** |
| **DNA17** | Sox2a | 25 bp | 1.159 | 23.348 | 0.785 | 0.215 | 4164013 | 3325000 |
| **DNA18** | Sox2a | 25 bp | 1.580 | 20.858 | 0.775 | 0.225 | 4831673 | 4395000 |
| **DNA19** | Sox2a | 25 bp | 0.586 | 15.580 | 0.841 | 0.159 | 4255561 | 5086266 |
| | | ***average*** | ***1.108*** | ***19.929*** | ***0.800*** | ***0.200*** | ***4417082*** | ***4268755*** |
| **DNA20** | Sox2a | 50 bp | 0.466 | 8.016 | 0.777 | 0.223 | 6912231 | 6720000 |
| **DNA21** | Sox2a | 50 bp | 0.693 | 8.245 | 0.518 | 0.482 | 8328949 | 6864202 |
| **DNA22** | Sox2a | 50 bp | 0.553 | 7.948 | 0.813 | 0.187 | 12388900 | 6075268 |
| | | ***average*** | ***0.571*** | ***8.070*** | ***0.703*** | ***0.297*** | ***9210027*** | ***6553156*** |
| **DNA23** | Sox2a | 100 bp | 1.093 | 10.345 | 0.534 | 0.466 | 12732300 | 8994022 |
| **DNA24** | Sox2a | 100 bp | 0.948 | 10.921 | 0.659 | 0.341 | 12375000 | 9024110 |
| **DNA25** | Sox2a | 100 bp | 0.994 | 14.942 | 0.873 | 0.127 | 17380000 | 10231400 |
| | | ***average*** | ***1.012*** | ***12.069*** | ***0.689*** | ***0.311*** | ***14162433*** | ***9416511*** |
| **DNA26** | Sox2a | 250 bp | 0.623 | 11.290 | 0.808 | 0.192 | 23067100 | 11868000 |
| **DNA27** | Sox2a | 250 bp | 0.889 | 7.570 | 0.621 | 0.379 | 16978200 | 12295000 |
| **DNA28** | Sox2a | 250 bp | 0.686 | 10.223 | 0.674 | 0.326 | 18811100 | 11363200 |
| | | ***average*** | ***0.733*** | ***9.694*** | ***0.701*** | ***0.299*** | ***19618800*** | ***11842067*** |
| **DNA29** | Sox2a | DNA- | 0.533 | 8.861 | 0.977 | 0.023 | 12393600 | 7241879 |
| **DNA30** | Sox2a | DNA- | 0.620 | 12.932 | 0.978 | 0.023 | 13535500 | 6436803 |
| **DNA31** | Sox2a | DNA- | 0.531 | 10.646 | 0.991 | 0.009 | 14978400 | 5911577 |
| | | ***average*** | ***0.561*** | ***10.813*** | ***0.982*** | ***0.018*** | ***13635833*** | ***6530086*** |

**Table S2. In vitro SMT analysis of Sox TFs on naked DNA**

#### Table S3

| TF | $k_a$ | | $K_R$ | | $s_L$ | |
|---|---|---|---|---|---|---|
| | Mean ($M^{-1}s^{-1}$) | SD | Mean | SD | Mean (bp) | SD |
| **Sox2** | 213372 | 12503 | 270.00 | 36.44 | 72.49 | 5.52 |
| **Sox2a** | 190747 | 24251 | 115.06 | 5.96 | 100.26 | 8.58 |
| **TF** | $p_{bind}$ | | $k_r$ | | $D_{1D}$ | |
| | Mean | SD | Mean ($s^{-1}$) | SD | Mean ($bp^2s^{-1}$) | SD |
| **Sox2** | 0.049 | 0.013 | 27000 | 3644 | 262769 | 39985 |
| **Sox2a** | 0.011 | 0.002 | 11506 | 596 | 502615 | 86043 |

**Table S3. Fit results of 1D target search model on specific on-rate**

#### Table S4

| Data label | TF | DNA | $t_1$ ($s^{-1}$) | $t_2$ ($s^{-1}$) | $t_3$ ($s^{-1}$) | $A_1$ | $A_2$ | $A_3$ | $k_{on}$ ($M^{-1}s^{-1}$) | $k_{s,on}$ ($M^{-1}s^{-1}$) |
|---|---|---|---|---|---|---|---|---|---|---|
| **NS01** | Sox2 | MN-6 | 0.647 | 3.071 | 8.940 | 0.686 | 0.241 | 0.073 | 23680500 | 8631691 |
| **NS02** | Sox2 | MN-6 | 0.882 | 3.454 | 8.981 | 0.725 | 0.195 | 0.079 | 24935700 | 7794559 |
| **NS03** | Sox2 | MN-6 | 0.846 | 3.805 | 10.072 | 0.612 | 0.246 | 0.142 | 26522700 | 10343600 |
| | | ***average*** | ***0.791*** | ***3.444*** | ***9.331*** | ***0.674*** | ***0.228*** | ***0.098*** | ***25046300*** | ***8923283*** |
| **NS04** | Sox2 | MN+2 | 1.158 | 5.254 | 10.383 | 0.554 | 0.305 | 0.140 | 29922600 | 10817400 |
| **NS05** | Sox2 | MN+2 | 0.976 | 5.713 | 15.098 | 0.417 | 0.410 | 0.172 | 21339600 | 10826200 |
| **NS06** | Sox2 | MN+2 | 0.875 | 4.723 | 13.196 | 0.366 | 0.335 | 0.299 | 23858800 | 11712200 |
| **NS07** | Sox2 | MN+2 | 0.404 | 3.724 | 15.026 | 0.358 | 0.393 | 0.249 | 20822500 | 11082700 |
| | | ***average*** | ***0.853*** | ***4.854*** | ***13.426*** | ***0.424*** | ***0.361*** | ***0.215*** | ***23985875*** | ***11109625*** |
| **NS08** | Sox2 | MN- | 0.529 | 1.906 | 49152 | 0.864 | 0.136 | 0.8824 | 13722200 | 3167949 |
| **NS09** | Sox2 | MN- | 0.650 | 2.800 | 50000 | 0.904 | 0.096 | 0.8521 | 13219000 | 3776514 |
| **NS10** | Sox2 | MN- | 0.743 | 2.126 | 50000 | 0.753 | 0.247 | 0.8782 | 11990700 | 3137987 |
| | | ***average*** | ***0.641*** | ***2.277*** | ***49717*** | ***0.840*** | ***0.160*** | ***0.871*** | ***12977300*** | ***3360817*** |
| **NS11** | Sox2 | CT+ | 0.580 | 2.209 | 8.777 | 0.869 | 0.082 | 0.049 | 70847900 | 11969600 |
| **NS12** | Sox2 | CT+ | 0.719 | 3.558 | 7.298 | 0.893 | 0.041 | 0.066 | 61060100 | 12995800 |
| **NS13** | Sox2 | CT+ | 0.669 | 3.862 | 12.608 | 0.701 | 0.217 | 0.082 | 67827900 | 13043700 |
| **NS14** | Sox2 | CT+ | 0.849 | 3.610 | 8.281 | 0.772 | 0.147 | 0.081 | 73940700 | 10640000 |
| | | ***average*** | ***0.704*** | ***3.310*** | ***9.241*** | ***0.809*** | ***0.122*** | ***0.069*** | ***68419150*** | ***12162275*** |
| **NS15** | Sox2 | CT- | 0.598 | 1.938 | 50000 | 0.928 | 0.072 | 0.8243 | 57719400 | 4403950 |
| **NS16** | Sox2 | CT- | 0.628 | 2.423 | 42535 | 0.935 | 0.064 | 0.8901 | 67695400 | 4905450 |
| **NS17** | Sox2 | CT- | 0.613 | 2.694 | 49984 | 0.837 | 0.163 | 0.9317 | 78568900 | 6181725 |
| | | ***average*** | ***0.613*** | ***2.351*** | ***47506*** | ***0.900*** | ***0.100*** | ***0.882*** | ***67994567*** | ***5163708*** |
| **NS18** | Sox2a | MN-6 | 0.595 | 3.546 | 11.634 | 0.711 | 0.225 | 0.063 | 9260000 | 5713612 |
| **NS19** | Sox2a | MN-6 | 0.620 | 2.198 | 13.112 | 0.850 | 0.112 | 0.038 | 12595000 | 6735666 |
| **NS20** | Sox2a | MN-6 | 0.391 | 3.840 | 9.170 | 0.758 | 0.163 | 0.078 | 6960000 | 6454677 |
| | | ***average*** | ***0.535*** | ***3.194*** | ***11.306*** | ***0.773*** | ***0.167*** | ***0.060*** | ***9605000*** | ***6301318*** |
| **NS21** | Sox2a | MN+2 | 0.346 | 2.336 | 14.979 | 0.415 | 0.367 | 0.218 | 5530231 | 3494397 |
| **NS22** | Sox2a | MN+2 | 0.475 | 4.008 | 18.330 | 0.379 | 0.495 | 0.125 | 6474098 | 3995655 |
| **NS23** | Sox2a | MN+2 | 0.768 | 5.218 | 17.754 | 0.392 | 0.313 | 0.295 | 6617896 | 4347224 |
| **NS24** | Sox2a | MN+2 | 0.293 | 2.426 | 13.782 | 0.334 | 0.457 | 0.210 | 8621147 | 4968547 |
| | | ***average*** | ***0.471*** | ***3.497*** | ***16.211*** | ***0.380*** | ***0.408*** | ***0.212*** | ***6810843*** | ***4201456*** |
| **NS25** | Sox2a | MN- | 0.426 | 2.211 | 49152 | 0.882 | 0.118 | 3E-05 | 7987725 | 5232614 |
| **NS26** | Sox2a | MN- | 0.463 | 2.145 | 50000 | 0.852 | 0.147 | 0.0005 | 8113179 | 3123943 |
| **NS27** | Sox2a | MN- | 0.693 | 2.683 | 50000 | 0.878 | 0.122 | 3E-31 | 9824681 | 3326255 |
| | | ***average*** | ***0.527*** | ***2.346*** | ***49717*** | ***0.871*** | ***0.129*** | ***0.000*** | ***8641861*** | ***3894271*** |
| **NS28** | Sox2a | CT+ | 0.449 | 3.907 | 14.459 | 0.909 | 0.076 | 0.015 | 10723200 | 6105000 |
| **NS29** | Sox2a | CT+ | 0.550 | 2.453 | 15.665 | 0.890 | 0.098 | 0.013 | 19436500 | 7133532 |
| **NS30** | Sox2a | CT+ | 0.556 | 4.055 | 15.498 | 0.886 | 0.105 | 0.009 | 24777800 | 7423283 |
| | | ***average*** | ***0.518*** | ***3.472*** | ***15.207*** | ***0.895*** | ***0.093*** | ***0.012*** | ***18312500*** | ***6887272*** |
| **NS31** | Sox2a | CT- | 0.444 | 3.569 | 50000 | 0.824 | 0.176 | 1E-31 | 23652300 | 4600628 |
| **NS32** | Sox2a | CT- | 0.382 | 2.797 | 50000 | 0.890 | 0.110 | 1E-15 | 16114500 | 5236463 |
| **NS33** | Sox2a | CT- | 0.624 | 5.696 | 50000 | 0.932 | 0.068 | 3E-20 | 19048200 | 5790161 |
| | | ***average*** | ***0.483*** | ***4.021*** | ***50000*** | ***0.882*** | ***0.118*** | ***0.000*** | ***19605000*** | ***5209084*** |

**Table S4. In vitro SMT analysis of Sox TFs on nucleosomes**
